# Ketogenic interventions restore cognition and modulate peripheral metabolic dysfunctions in Alzheimer’s disease mouse models

**DOI:** 10.1101/2024.09.04.610637

**Authors:** Paule E. H. M’Bra, Laura K. Hamilton, Gaël Moquin-Beaudry, Myriam Aubin, Chenicka L. Mangahas, Federico Pratesi, Elsa Brunet-Ratnasingham, Anne Castonguay, Sophia Mailloux, Manon Galoppin, Megan Bernier, Marta Turri, Annick Vachon, Marian Mayhue, Anne Aumont, Martine Tétreault, Mélanie Plourde, Karl J.L. Fernandes

## Abstract

Lifestyle factors modulate dementia risk. We investigated mechanisms of dementia risk reduction by emerging dietary ketogenic interventions. We show that distinct interventions, a medium-chain triglycerides (MCT)-enriched diet and a carbohydrate-free, high-fat diet (CFHF), improve cognition and dendritic spine density of memory-associated hippocampal neurons in two mouse models of Alzheimer’s disease (AD). Only the CFHF diet drove increased circulating ketones, suggesting distinct underlying mechanisms. AD mice exhibited baseline and diet-induced susceptibility to peripheral metabolic disturbances that were improved by MCT and exacerbated by CFHF diets. Prominent AD-associated dysregulation of the liver transcriptome was largely restored by both interventions, but MCT also downregulated lipogenic enzymes and did not trigger a CFHF-like inflammatory signature. Novel AD- and diet-induced plasmatic changes in hormones and lipid species were identified. Thus, different ketogenic interventions yield cognitive benefits in AD models while showing intervention-specific modulation of peripheral metabolic defects, with implications for design of therapeutic ketogenic strategies.

## INTRODUCTION

Alzheimer’s disease (AD), the leading cause of dementia, is characterized by a progressive decline of learning, memory and other cognitive functions that compromise the ability to perform daily tasks^1^. Symptom severity increases along the AD continuum from mild cognitive impairment (MCI) to severe dementia^2^ and is paralleled by the progression of brain changes which start even before MCI^3,4^. These brain changes include extracellular aggregates of amyloid plaques, intracellular neurofibrillary tangles, neuroinflammation, synaptic and neuronal loss, cerebral atrophy especially in the hippocampus and impaired glucose metabolism^2,5–7^. Rare cases of familial AD are caused by inherited autosomal dominant genetic variations coding for amyloid precursor protein (APP), presenilin 1 and 2^8^. Most cases of AD are sporadic with primary non-modifiable risk factors that include aging and genetic predispositions (such as apoE4 allele^2,9,10^), as well as modifiable lifestyle risk factors that contribute up to 40% of disease risk^1,5,11^. Factors influencing peripheral metabolism, such as type 2 diabetes (T2D) and obesity, are significant promoters of AD risk, and recent studies have highlighted the implication of peripheral dysfunctions in AD pathophysiology^12–16^.

Ketogenic interventions (KI) are emerging as promising strategies for treating patients with AD. They are based on the premise that the brain’s impaired glucose metabolism in AD^3,4,6,17–19^ can be compensated for by increasing circulating levels of ketone bodies, an energy source for the brain whose use is preserved in aging and AD^17,20^. Two common KI include adherence to a low-carbohydrate/high-fat diet, which triggers a metabolic switch that boosts hepatic ketone production through increased oxidation of endogenous fatty acids^21^, or dietary supplementation with medium-chain triglycerides (MCT)^22^, which are likewise metabolized by the liver to produce ketones. The length of the medium-chain fatty acids used during MCT supplementation influences the ability to produce ketones, with capric acid (C10:0) and especially caprylic acid (C8:0) having stronger ketogenic effects than lauric acid (C12:0)^23–25^. Low carbohydrate and MCT diets have both been reported to yield cognitive benefits in AD or MCI^26^. For example, a 6-month dietary supplementation with ketogenic MCTs improved memory in patients with MCI^23^. AD patients who adhered to a ketogenic low-carbohydrate/high-fat diet for 12 weeks presented improvements of their daily functions and quality of life^27^.

Design of optimized, next-generation KI requires deeper mechanistic understanding of the factors determining their efficacy. For example, some patients fail to exhibit cognitive benefits despite increased plasma ketone levels^23^ and the therapeutic time window for optimal effects is not clear. Ketones also have understudied non-energetic roles^28^ and ketogenic substrates such as MCT can potentially have direct, ketone-independent effects that are largely unexplored. For instance, research in epilepsy suggests that MCTs can directly cross the blood-brain barrier to influence neurotransmission in the brain^22,29^. Given the relationship between AD and peripheral metabolic disorders, it is also important to consider the impact of KI on peripheral metabolism and whether genetic predisposition to AD affects the ability to respond to such interventions^30,31^. Here, we modeled MCT and low carbohydrate high-fat KI in transgenic mouse models of AD, investigating whether they protect against cognitive and anatomical decline in the brain and how they affect key measures of peripheral metabolism.

## RESULTS

### MCT and CFHF diets reverse cognitive deficits in AD mice and increase spine density in pre-symptomatic AD mice

We first asked whether the cognitive benefits of KI in AD can be reproduced using AD mouse models. To do so, we tested whether KI can reverse the learning and memory losses seen in symptomatic 5xFAD mice, which exhibit high amyloid plaque load by 2 months of age and reproducibly developing cognitive impairments by 5 months of age ^32^.

5xFAD mice at 6 months old were placed on one of three diets for one month: either a Control diet (70% carbohydrates, 20% fats and 10% proteins), an MCT-enriched diet in which half of the fat content consists of ketogenic MCTs (C8:0/C10:0 ratio of 3:2), or a carbohydrate-free, high-fat diet (CFHF) (**Fig 1a-b, Supp. tables 1-2)**. After one month on these diets, learning and memory were assessed using the Morris Water Maze. Learning, defined as a decrease of escape latency during the 4-days training phase, was significantly slower in 5xFAD mice versus their wildtype (WT) littermates and was significantly improved by the MCT diet (**Fig 1c-d)**. Spatial memory, gauged by preference for the target quadrant versus the opposite quadrant upon platform removal, was significantly impaired in 5xFAD mice compared to WT mice, and was restored by both the MCT and CFHF diets (**Fig 1e-f**). Thus, one month of KI was sufficient to reverse learning and memory deficits in symptomatic AD mice.

**Figure 1:**
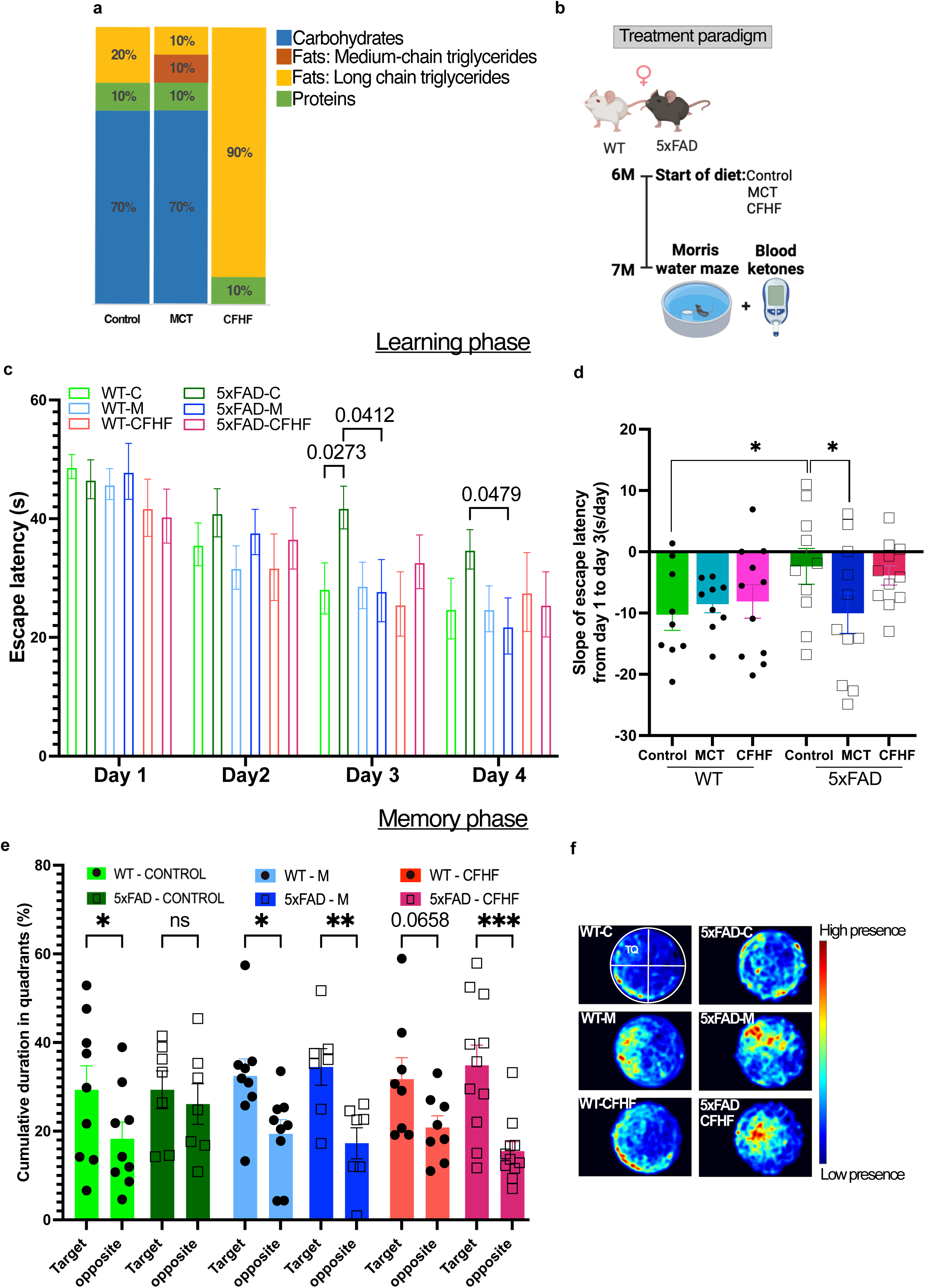
Both KI reverse learning and memory deficits in AD mice. a) Macronutrients proportion in the three diets (Details in Supplementary tables 1 – 2). b-f) Morris water maze results following ketogenic interventions (KI) in symptomatic 5xFAD mice: b)Timeline of dietary interventions to reverse cognitive decline (n=7-11 animals/genotype/diet), c) Average escape latency of 4 trials/day (Mixed effects model showing time effect (p-value <0.0001, F(2.747,141.9)=27.67) and Fisher post hoc test(detailed on graph)), d)Average slope of each mouse learning curve from day 1 to day 3 (2-way ANOVA showing a trend in genotype factor (p-value = 0.098, F(1,55) = 2.857) and Fisher post hoc test (detailed on graph)), e) Ability to discriminate target quadrant from opposite quadrant assessed by the cumulative percentage time spent in quadrants dring memory phase (2-way ANOVA showing significant quadrant discrimination factor (p-value <0.0001, F(1,90)=28.19) and Fisher post hoc test comparing Target vs Opposite quadrant for each group (detailed on graph)). f) Merged heatmaps per group indicating the preferred location of the mice. The pool is virtually separated in 4 quadrants : the target quadrant (TQ) contained the platform during the training phase Fisher post hoc test: *p-value<0.05; **p-value<0.01; ***p-value<0.001; ****p-value<0.0001

AD-associated learning and memory deficits are the outcome of brain pathologies that begin accumulating during presymptomatic stages of the disease. We therefore investigated whether KI administered at pre-symptomatic stages can prevent/delay loss of dendritic spines in two separate mouse models of AD, namely 5xFAD and 3xTg-AD. While the rapidly progressing 5xFAD model simulates late stages of AD^32^, the 3xTg-AD model is a slowly progressing model that facilitates study of early stages of AD^33^. Because of the difference in disease progression between the models, 5xFAD mice were administered diets for three months (from 2- to 5-months-old) and 3xTg-AD mice for five months (from 3– to 8-months-old) (Fig 2a). Golgi-cox staining was performed to label the dendrites and the density of dendritic spines was quantified on two defined neuronal populations of the spatial learning- and memory-associated dorsal hippocampus ^34,35^, namely, dentate gyrus (DG) granule neurons and CA1 pyramidal neurons (**Fig 2b-e)**. Quantification showed that baseline changes in the density of dendritic spines were not yet present on these neuronal populations at these ages in either the 5xFAD or 3xTg-AD models (**Control diet, Fig 2f-m**). Interestingly, the MCT and CFHF interventions stimulated region-specific increases in dendritic spines in both WT strain controls and AD mouse models. In the granule neurons of the DG, MCT increased dendritic spines in 3° dendrites of 5xFAD mice and both 2° and 3° dendrites of their WT strain (**Fig 2f-g**), as well as in 2° and 3° dendrites of 3xTg-AD mice (**Fig 2j-k**). Similarly, CFHF diet increased dendritic spine density of 3° dendrites in 5xFAD mice and in 2° and 3° dendrites of their WT strain controls (**Fig 2f-g**). Neither MCT or CFHF diets stimulated significant increases in spine density of CA1 pyramidal neurons in either the 5xFAD, 3xTg-AD or their respective strain controls (**Fig 2h-i, l-m**).

**Figure 2:**
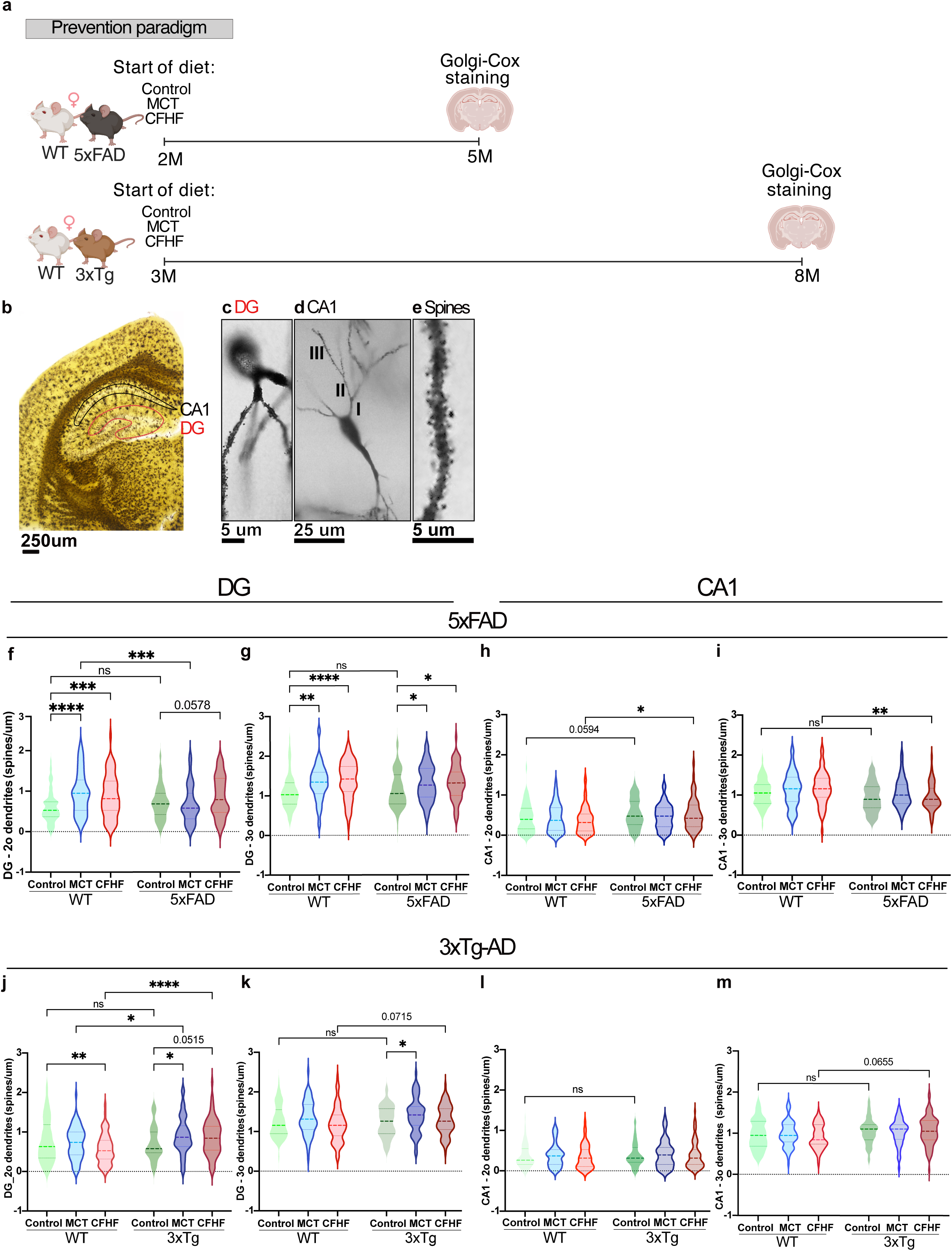
Both KI increase the dendritic spine density in pre-symptomatic AD mice. a) Timeline of 3 months in 5xFAD mice (n=6 animals/genotype/diet) and 5 months in 3xTg-AD mice (n=6 animals/strain/diet). b-e) Representative pictures of neurons stained with Golgi-Cox staining in a brain section (b, 4x magnification, scale: 25um) and within the regions of interest : c) the dentate gyrus (DG, 100x magnification, scale: 5um) and d) the cornu ammonu 1 (CA1, 100x magnification, scale: 25um). The spines (e, scale: 5um) were quantified on secondary (2o,represented by **II** on picture) and tertiary (3o,represented by **III** on picture) dendrites (n=50-90 neurons/strain/diet). f-g) Quantification of spine density within the DG of 5xFAD mice. f) 2o dendrites (2-way ANOVA showing diet effect (p-value=0.0003, F(2,447)=8.178) and interaction between diet and genotype (p-value=0.0008, F(2,447)=7.310).Fisher post hoc test (detailed on graph) ). g)3o dendrites (2-way ANOVA showing diet effect (p-value<0.0001, F(2,446)=12.91).Fisher post hoc test (detailed on graph)). j-k) Quantification of spine density within the DG of 3xTg-AD mice. j) 2o dendrites (2-way ANOVA showing strain effect (p-value=0.0010, F(1,442)=11.07) and interaction between diet and strain (p-value=0.0038, F(2,442)=5.657). Fisher post hoc test (detailed on graph)). k) 3o dendrites (2-way ANOVA showing diet effect (p-value=0.0060, F(2,440)=5.180) and a trend in strain factor (p-value=0.069, F(1,440)=3.314). Fisher post hoc test (detailed on graph)). h-i) Quantification of spine density within the CA1 of 5xFAD mice. h) 2o dendrites (2-way ANOVA showing genotype effect (p-value=0.0040, F(1,390)=8.399).Fisher post hoc test (detailed on graph)). i)3o dendrites(2-way ANOVA showing genotype effect (p-value=0.0001, F(1,390)=14.71).Fisher post hoc test (detailed on graph)). l-m) Quantification of spine density within the CA1 of 3xTg-AD mice. l) 2o dendrites(2-way ANOVA : ns). m) 3o dendrites(2-way ANOVA showing strain effect (p-value=0.0031, F(1,405) =8.875). Fisher post hoc test (detailed on graph)) Fisher post hoc test: *p-value<0.05; **p-value<0.01; ***p-value<0.001; ****p-value<0.0001

These data reveal that KI reverse learning and memory defects in AD mouse models and increase dendritic spine density in sub-populations of hippocampal neurons even at presymptomatic stages of the disease.

#### Cognitive benefits of MCT supplementation do not rely on permanently elevated blood ketones

Ketogenic diets are thought to increase brain energy uptake in AD by elevating levels of circulating ketones^18^, most notably beta-hydroxybutyrate (BHB), and by reducing or stabilizing blood glucose levels, together resulting in a lower glucose/ketone index (GKI<10)^36^. Unexpectedly, despite the similar effects of MCT and CFHF on cognition and dendritic spines, analysis of blood ketones and glucose levels in the above cohort of 5xFAD mice suggested important differences between these dietary interventions. Only CFHF triggered elevated blood BHB levels, which were present at both the 14-day and 35-day timepoints after start of the diet **(Supp.Fig 1a-b).** Even so, MCT and CFHF were both able to gradually reduce blood glucose levels **(Supp.Fig 1b).**

To explore this observation in greater depth, we performed 6-month MCT and CFHF interventions in a new cohort of 3xTg-AD mice, whose slower time-course of pathologies facilitates temporal analysis of disease progression **(Fig 3a).** As was seen in 5xFAD mice, only CFHF elevated BHB levels in 3xTg-AD mice **(Fig 3b-c, Supp.Fig 1a)**, with increased BHB clearly detectable at the 14-day timepoint. Interestingly, 3xTg-AD mice displayed a ketone increase in response to the CFHF diet that was dampened compared to their WT strain (**Fig 3b – Short-term, Long-term**). 3xTg-AD mice also had elevated blood glucose levels that were present even before dietary interventions started (**Fig 3c – Baseline**) and that were further increased by the CFHF diet (**Fig 3c – Short-term, Long-term**). Calculation of the GKI indicated that the dampened ketone elevation and high glucose levels in 3xTg-AD mice on CFHF diet resulted in only a mild ketosis state (6 <GKI<9) at day 14 and a non-ketosis state at day 171 (GKI > 9) **(Supp.Fig 1d** – **Short-term, Long-term**). While MCT again did not boost blood ketone levels, and it still tended to reduce blood glucose levels compared to Control diet **(Fig 3c, Supp.Fig 1c).**

**Figure 3:**
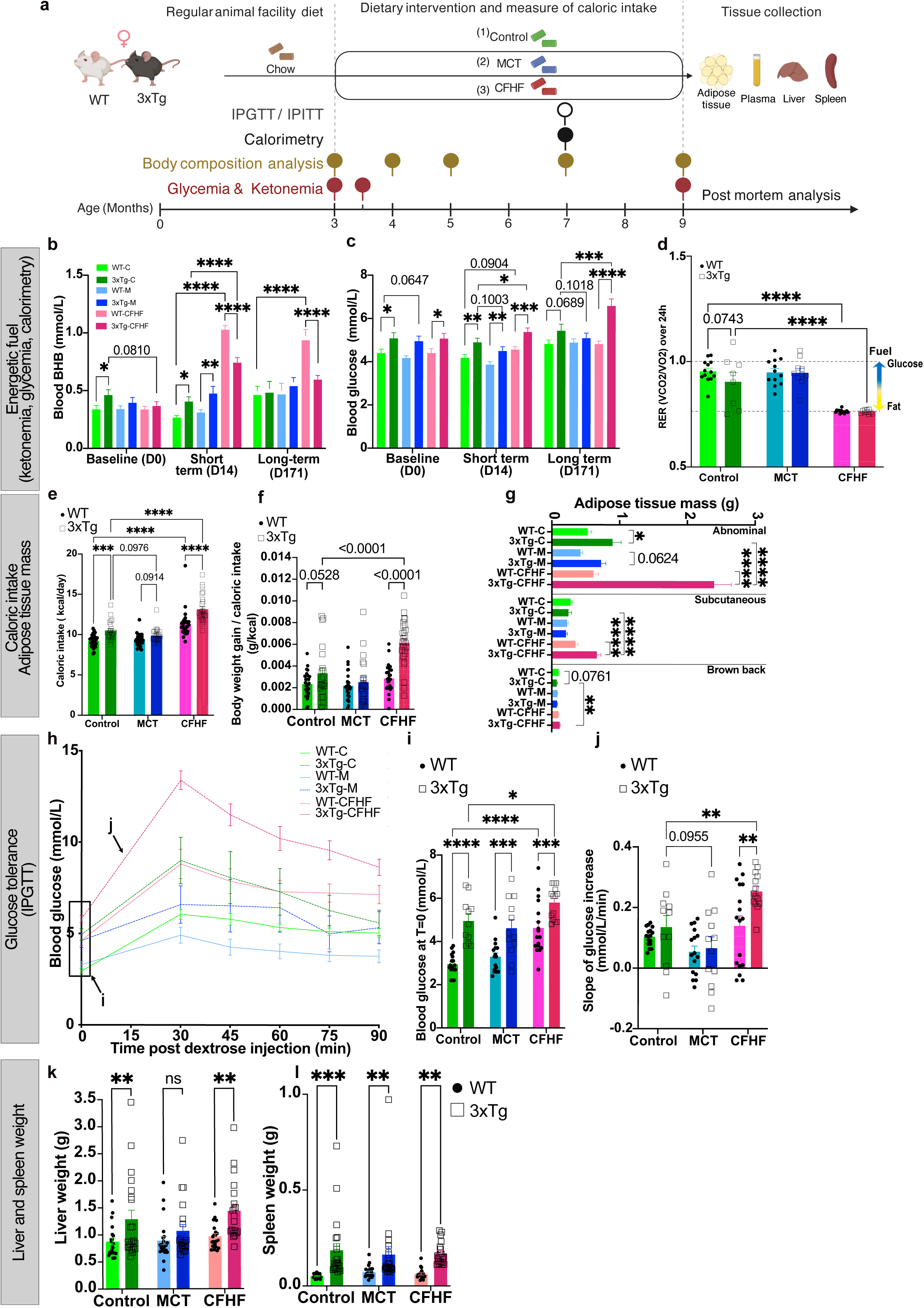
Physiological response of pre-symptomatic 3xTg-AD mice to ketogenic diets. a) Timeline: 3 months old female 3xTg-AD and their WT strain mice, received either a Control diet, a MCT-enriched diet or a CFHF diet for 6 months alongside with metabolic measures: glycemia and ketonemia, body composition analysis, indirect calorimetry and intraperitoneal glucose/insulin tolerance test (IPGTT, IPITT). Plasma and some peripheral tissues were collected for post-mortem analysis. b-d) Evaluation of energetic fuel source : b-c) Longitudinal monitoring of glycemia and ketonemia in 3xTg-AD mice (n=17-21/strain/diet) at Baseline(day 0), Short-term (day 14) and Long-term (day 171) timepoints. b) Blood ketones levels (2-way ANOVA for each timepoint (<u>Baseline</u>: strain effect (p-value=0.0234, F(1,108)=5.291) with no effect of diet consistent with the fact that mice were on the same diet at this level, <u>Short-term</u>: diet effect (p-value<0.0001, F(2,115)=117.3) and interaction between diet and strain (p-value<0.0001, F(2,115)=20.47, <u>Long-term</u>: diet(p-value<0.0001, F(2,112)=20.54) and strain effects (p-value=0.0480, F(1,112)=3.997) with interaction (p-value<0.0001, F(2,112)=10.07). Fisher post hoc test (detailed on graph)). c) Blood glucose levels(2-way ANOVA for each timepoint (<u>Baseline</u>: strain effect(p-value<0.0001, F(1,114)=17.31) with no effect of diet consistent with the fact that mice were on the same diet at this level, <u>Short-term</u>: diet(p-value<0.0001, F(2,114)=11.66) and strain effects(p-value<0.0001, F(1,114)=28.55 with no interaction, <u>Long-term</u>: diet (p-value=0.0051, F(2,112)=5.533) and strain effects(p-value<0.0001, F(1,112)=20.98) with interaction(p-value<0.0024, F(2,112)=6.368). Fisher post hoc test (detailed on graph)). d) Respiratory exchange ratio(RER) average of 24h calorimetric measures (n=8-12/strain/diet, 2-way ANOVA showing diet effect (p-value<0.0001, F(2,60)=62.43). Fisher post hoc test (detailed on graph)). e-g) Energy intake and body fat : e) Average daily food intake over 6 months on diet (n=22-28/strain/diet, 2-way ANOVA showing diet(p-value<0.0001, F(2,148)=71.68) and strain effects (p-value<0.0001, F(1,148)=39.57) with interaction(p-value=0.0314, F(2,148)=3.544). Fisher post hoc test (detailed on graph)). f) Body weight gain per caloric intake (2-way ANOVA showing diet(p-value<0.0001, F(2,148)=21.83) and strain effects (p-value<0.0001, F(1,148)=29.54) with interaction (p-value<0.0001, F(2,148)=10.36). Fisher post hoc test (detailed on graph)). g) Fat pad dissection (n=17-24/strain/diet): Abdominal fat (2-way ANOVA showing diet(p-value<0.0001, F(2,124)=37.39) and strain(p-value<0.0001, F(1,124)=72.76) effects with interaction (p-value<0.0001, F(2,124)=25.70). Fisher post hoc test (detailed on graph)), Subcutaneous fat (2-way ANOVA showing significant diet(p-value<0.0001, F(2,125)=35.02) and strain(p-value=0.0046, F(1,125)=8.319) effects with interaction(p-value<0.0001, F(2,125)=13.50). Fisher post hoc test (detailed on graph)), Brown back fat (2-way ANOVA showing diet effect (p-value<0.0184, F(2,125)=4.127) and trend toward interaction between strain and diet(p-value=0.0775, F(2,125)=2.611).Fisher post hoc test (detailed on graph)) h-j) Intra peritoneal glucose tolerance test (IPGTT) after 4 months on diet (n=11-17/strain/diet) showing : h) Curves of blood glucose change after IP injection of glucose (Mixed-effects model showing time (p-value<0.0001, F(2.477, 199.6)=62.92) and experimental group (p-value<0.0001, F(5, 81)=16.30) effects with interaction (p-value<0.0001, F(25, 403)=4.008). i)fasting glucose levels (2-way ANOVA showing diet(p-value<0.0001, F(2,81)=17.25) and strain(p-value<0.0001, F(1,81)=52.60) effects. Fisher post hoc test (detailed on graph)). j)Rate of glucose increase 30 min after injection (2-way ANOVA showing diet(p-value<0.0001, F(2,81)=14.07) and strain(p-value=0.017, F(1,81)=5.941) effects.Fisher post hoc test (detailed on graph)). k-l) After 6 months on diet, liver and spleen were examined k)Liver weight (n=20- 22/strain/diet, 2-way ANOVA showing strain effect (p-value<0.0001, F(1,120)=16.57). Fisher post hoc test (detailed on graph)). l)Spleen weight (n=19-22/strain/diet, 2-way ANOVA showing strain effect (p-value<0.0001, F(1,119)=33.16). Fisher post hoc test (detailed on graph)) Fisher post hoc test: *p-value<0.05; **p-value<0.01; ***p-value<0.001; ****p-value<0.0001

We confirmed the resting metabolic state of 3xTg-AD mice by measuring the volume of respiratory gases during 24h in calorimetric cages. The respiratory exchange ratio (RER) is the ratio of produced carbon dioxide (VCO2) out of consumed dioxygen (VO2). Mice on both the Control and MCT diets had a RER near 1 meaning a higher glucose energy contribution and therefore were not in ketosis **(Fig 3d, Supp.Fig 1e)**. In contrast, the CFHF fed mice had a RER of 0.7 indicating that they primarily utilized fat as their energy substrate which suggest a ketosis state (**Fig 3d, Supp.Fig 1e)**. The circadian rhythms affected the RER of MCT and Control fed mice which tended to be lower during light cycle. In the mice fed with CFHF, we did not observe this variation indicating a permanent ketosis **(Supp.Fig 1e).**

Collectively, these data reveal that CFHF, but not MCT, drives a permanent ketosis through a metabolic shift towards fat oxidation. 3xTg-AD mice have a dampened ketogenic response to the CFHF diet and display signs of hyperglycemia, while 3xTg-AD mice on MCT diet tend towards a normalization of their blood glucose levels. These data indicate that in the case of the MCT diet, neurological improvements in the brain do not correlate with sustained increase in blood ketones.

### MCT and CFHF diets in pre-symptomatic AD mice have opposing effects on AD- induced peripheral defects

The differential glycemia and dampened ketone response observed in 3xTg-AD mice raised the question of whether genetic risk for AD can trigger peripheral metabolic defects and whether KI could attenuate these defects. We explored this question using pre-symptomatic 3xTg-AD mice, which we found exhibited significant peripheral changes under basal dietary conditions **(Supp.Fig 2)**. This included increased body weight already at 2- months old which became more prominent with age **(Supp.Fig 2a-b)**. They also developed increased adiposity, especially white abdominal fat as observed by CT scan and by fat pad dissections **(Supp.Fig 2c-f)**, elevated resting blood glucose **(Supp.Fig 2k)**, and hepatosplenomegaly **(Supp.Fig 2g-j)**. These alterations are suggestive of disease-induced systemic metabolic aberrations.

We then asked whether these systemic alterations were affected in mice subjected to KI. While 3xTg-AD mice on the Control diet again exhibited significantly higher caloric intake, body weight gain and abdominal fat than WT strain controls (**Fig 3e-g**), these differences were all reduced in 3xTg-AD mice on the MCT diet and strikingly increased in 3xTg-AD mice on the CFHF diet (**Fig 3e-g**). CFHF diet significantly and selectively increased weight gain in 3xTg-AD mice, doubling the weight gain compared to the Control diet, while it did not notably impact the body weight of WT mice (**Fig 3f**).

Longitudinal body composition analysis with EchoMRI corroborated that 3xTg-AD mice had higher fat mass (**Supp. Table 3**). CFHF diet increased the alterations of the body composition in 3xTg-AD mice, including higher fat mass percentage, lower lean mass, and reduced water percentages while it impacted these parameters minimally or not at all in WT mice (**Supp. Table 3: 3xTg-C vs 3xTg-CFHF, WT-C vs WT-CFHF**). Conversely, 3xTg-AD mice on MCT diet exhibited a body composition closer to that of WT mice from D112 (**Supp. Table 3: WT-M vs 3xTg-M**). Consistent with EchoMRI, fat pad dissections revealed that 3xTg-AD mice on CFHF presented a threefold increase of abdominal fat weight compared to 3xTg-AD mice on Control diet (**Fig 3g**). The difference between 3xTg-AD mice and WT mice increased from 1.6-fold on Control diet to fourfold on CFHF, showing the selective impact of CFHF on abdominal fat in 3xTg-AD mice **(Fig 3g)**. These results show that 3xTg-AD mice presented higher baseline food intake, body weight and body fat, especially abdominal fat. MCT alleviated these features while CFHF selectively exacerbated them in 3xTg-AD and not in WT mice.

Since 3xTg-AD mice showed signs of hyperglycemia (**Fig 3c, Supp.Fig 1c, Supp.Fig 2k**), we evaluated glucose clearance and insulin sensitivity with intraperitoneal glucose (IPGTT) and insulin tolerance tests (IPITT) after 4 months on diet (**Fig 3h-j, Supp.Fig 1f-k)**. IPGTT and IPITT confirmed hyperglycemia, showing higher fasting glucose in 3xTg-AD mice on Control diet **(Fig 3i –Control**, **Supp.Fig1h – Control)**. 3xTg-AD mice also exhibited impaired glucose handling as reflected by higher glucose levels in response to glucose or insulin injection (**Fig 3h, Supp.Fig 1k**). The administration of MCT diet enhanced glucose tolerance, as evidenced by a reduction in overall blood glucose levels and improved initial response to glucose injection (**Fig 3h, 3j**). In contrast, CFHF diet increased plasma glucose in both WT and 3xTg-AD mice when compared to Control diet fed mice (**Fig 3h-i**). 3xTg-AD mice on CFHF exhibited a more pronounced hyperglycemia and a more abrupt initial response to glucose load compared to WT mice on CFHF and 3xTg-AD mice on Control diet (Fig2i-j). Surprisingly, the response of 3xTg-AD mice to insulin was unchanged by KI, as evidenced by an unaltered slope of the insulin-induced hypoglycemia (**Supp.Fig 1i**). However, the effect of insulin appeared to be shorter in CFHF fed mice, as blood glucose levels at T=90min compared to T=0 were 14% higher in CFHF diet fed mice, while it was around 17% decreased in Control diet fed mice **(Supp.Fig 1g, 1j)**. Altogether, 3xTg-AD mice presented impaired glucose tolerance, yet without insulin resistance. MCT reduced the impaired glucose metabolism while CFHF accentuated it. WT mice also developed hyperglycemia on CFHF, but less than 3xTg-AD mice on the same diet.

3xTg-AD mice exhibited hepatosplenomegaly with their spleen weight being almost fourfold that of WT mice on the same diet (**Fig 3k** –Liver weight: WT-Control = 0.88±0.06g; 3xTg-Control = 1.29±0.17g; **Fig 3l -** Spleen weight: WT-Control = 0.05±0.003g; 3xTg-Control = 0.186±0.04g). CFHF did not affect these differences<u>, but 3xTg mice on MCT tended to</u> <u>resemble WT mice</u> (**Fig 3k** –Liver weight: 3xTg-MCT = 1.07±0.12g; WT-Control = 0.88±0.06g, p=0.20).

Collectively, these results show that AD-induced peripheral metabolic disturbances are enhanced when combined with CFHF diet, while MCT diet mitigates them.

### AD-induced changes in the liver are prevented by KI

The liver is the first organ through which absorbed dietary nutrients pass after leaving the gut and it plays a central role in the synthesis, modification, and storage of glucose, ketones, and fatty acids. Since 3xTg-AD mice presented signs of alterations in processing these substrates and exhibited hepatosplenomegaly, we investigated liver cytoarchitecture, gene expression and its response to KI.

Liver staining by Hematoxylin Eosin and neutral lipid staining with Oil Red O and BODIPY stains revealed that lipid droplets surface area were increased in livers of 3xTg-AD mice compared to WT mice (**Fig 4a-b**). This neutral lipid accumulation was unchanged by the MCT diet but was significantly reduced by CFHF diet in both WT and 3xTg-AD mice, remaining higher in the 3xTg-AD mice (**Fig 4a-b**).

**Figure 4:**
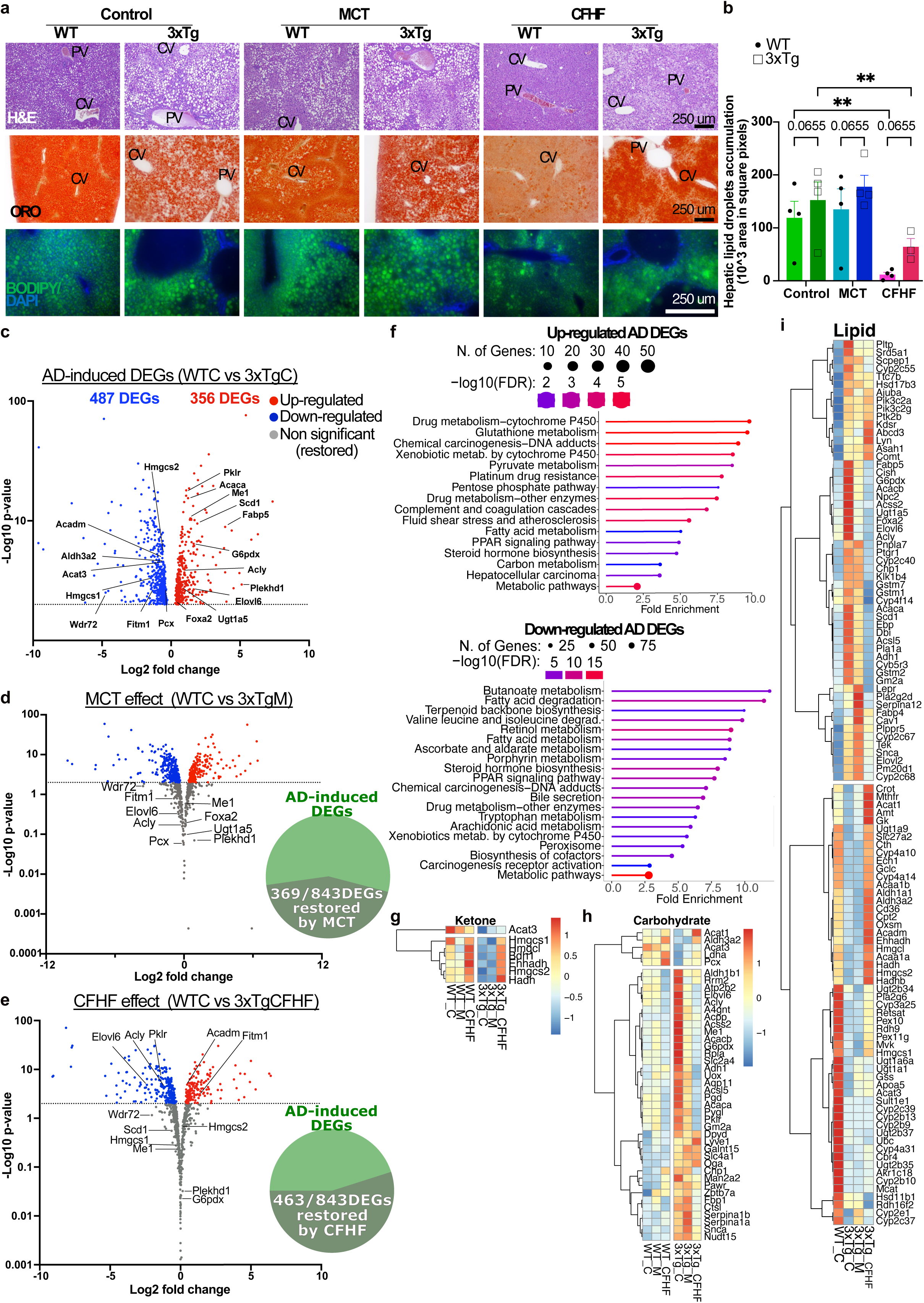
AD-induced changes in the liver are prevented by KI. a-b)Hepatic lipid content : a) Hepatocellular lipid droplet accumulation observed with hematoxylin-eosin (HE, magnification 10x), oil red o (ORO, magnification 10x) and BODIPY staining (magnification 20x), CV: central vein, PV: portal vein, scale bars:250um b) Quantification of lipid vacuoles from HE images with ImageJ (n=4/strain/diet, 2-way ANOVA showing diet effect(p-value=0.0007, F(2,19)=10.92) and strain factor nearly significant (p-value=0.0655, F(1,19)=3.821)). Fisher post hoc test: *p-value<0.05; **p-value<0.01; ***p-value<0.001; ****p-value<0.0001 c-i)Liver RNA-sequencing: c-e) Volcano plot of DEGs: d) between WT-C and 3xTg-C, e)between WT-C and 3xTg-M, f)between WT-C and 3xTg-CFHF. The volcano plots in 3e and 3f only show DEGs unaltered by diets (common DEGs between WT-C vs 3xTg-C and WT-C vs 3xTg-M (3e), common DEGs between WT-C vs 3xTg-C and WT-C vs 3xTg-CFHF (3f)) and genes restored by diets (significant DEGs in WT-C vs 3xTg-C, but not different in WT-C vs 3xTg-M or in WT-C vs 3xTg-CFHF) (n=4/strain/diet, DESeq2, cut-off p-value<0.01). Some genes were labelled to illustrate the restoration by diet (complete list of DEGs in Supp. table 4) The pie chart show the proportion of genes restored in AD-induced DEGs f) Top 20 pathways involving Up- and Down-regulated genes in 3xTg-C (KEGG, cut-off FDR<0.05) g-i) Heatmaps of DEGs regrouped by categories: h)Ketones, i)Carbohydrate, j)Lipid

To better understand these disease- and diet- effects on the liver we performed liver RNA-sequencing. On the Control diet, WT and 3xTg-AD mice exhibited 843 differentially expressed genes (DEGs, p<0.01), with a range of ten-fold upregulated to ten-fold downregulated (**Fig 4c**, Supp.Fig 3a-b, **Supp.Table 4**). Remarkably, 369/843 of these AD-induced DEGs were no longer significant in 3xTg-AD mice on the MCT diet and 463/843 in 3xTg-AD mice on the CFHF diet (**Fig 4d-e, Supp.Fig 3a-b**), suggesting a restoration of their expression level towards the level of WT mice on Control diet. MCT and CFHF diets also stimulated 731 and 1313 “new” changes in gene expression, respectively (**Supp.Fig 3a**, **Fig 5**).

**Figure 5:**
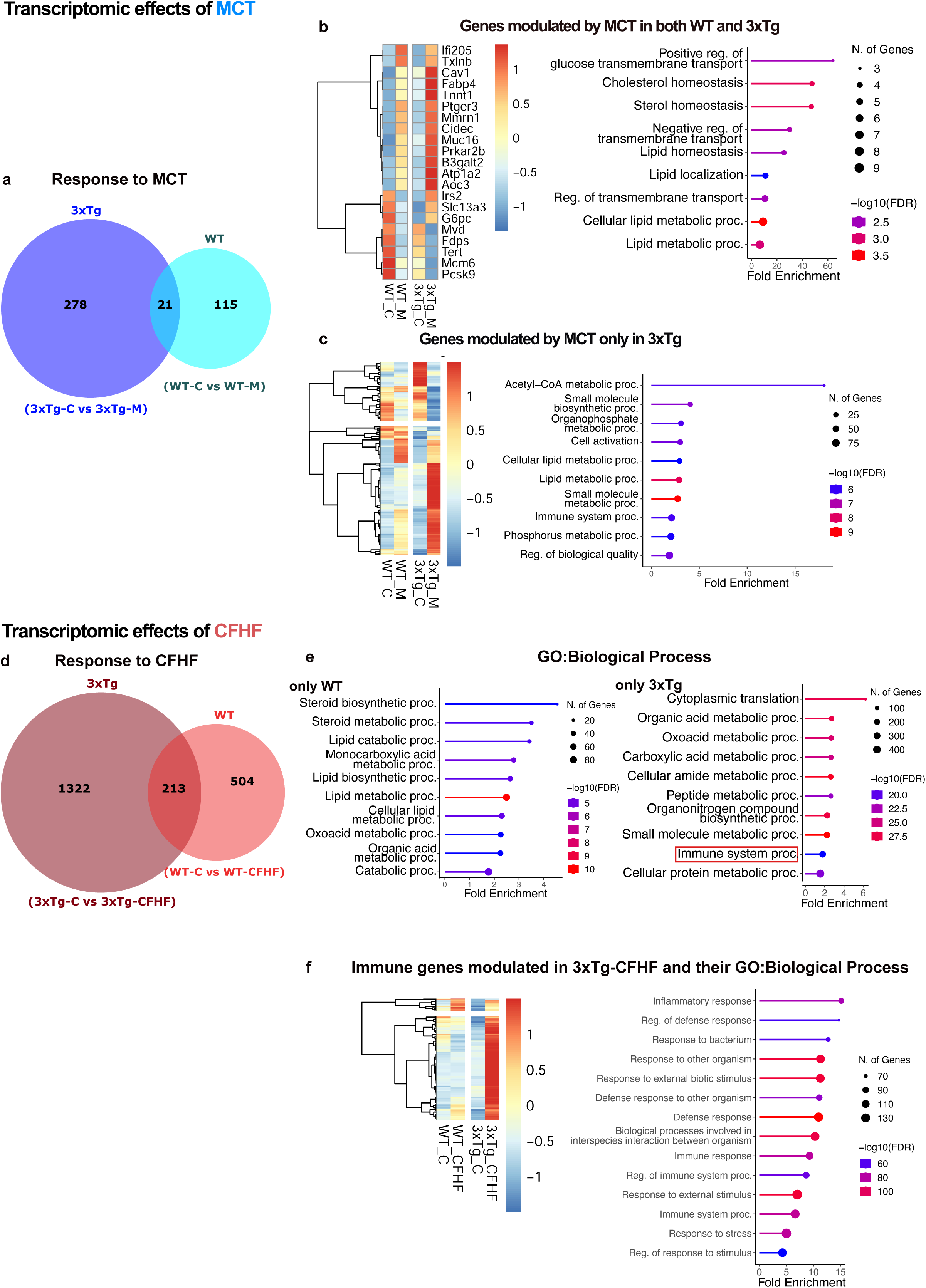
MCT and CFHF triggered second class transcriptomic changes in the liver of AD mice. Second class DEGs are genes modulated by KI and not modulated by AD mutations. a-c) Transcriptomic effects of MCT : a) Venn diagram of DEGs comparing the response to MCT of 3xTg-AD vs WT mice : 3xTg = 3xTg-C vs 3xTg-M ; WT = WT-C vs WT-M. b) Heatmap of genes significantly modulated in both 3xTg-AD and WT mice fed with MCT diet compared to their counterpart on Control diet (group mean, n=4/strain/diet, DESeq2, cutoff p-value<0.01), and Top 10 pathways associated to these genes (GO Biological Process, FDR<0.05). c) Heatmap of genes exclusively modulated in 3xTg-AD mice on MCT diet (group mean, n=4/strain/diet, DESeq2, cutoff p-value<0.01) with associated Top 10 pathways(GO Biological Process, FDR<0.05) d-g) Transcriptomic changes induced by CFHF : d)Venn diagram of DEGs comparing the response to CFHF of 3xTg-AD vs WT mice: 3xTg = 3xTg-C vs 3xTg-CFHF ; WT = WT-C vs WT-CFHF. e)Top 10 GO Biological Processes involving DEGs in the following contrasts : WT-C vs WT-CFHF and 3xTg-AD vs 3xTg-CFHF and highlighting “Immune system process” as a difference between the two contrasts(cutoff FDR<0.05). f)Heatmap of CFHF-induced upregulated immune genes in 3xTg-AD mice (n=4/strain/diet, DESeq2, cutoff p-value<0.01) and related Top 15 GO Biological Processes associated (FDR<0.05).

AD-induced pathway changes were identified by using AD-induced DEGs for Gene set enrichment analysis based on Gene Ontology biological process (GO: BP), Gene Ontology cellular component (GO:CC) and Kyoto Encyclopedia of Genes and Genome (KEGG) analyses. The AD-induced pathway changes were regrouped into 5 categories, namely “Lipids” (13% of AD-induced DEGs, 110/843), “Ketones” (0.8% of AD-induced DEGs, 7/843), “Carbohydrate-derivative processes” (5% of AD-induced DEGs, 42/843), “Mitochondrial structure” (7.9% of AD-induced DEGs, 67/843), and “Xenobiotic processes” (5.7% of AD-induced DEGs, 48/843)(**Fig 4f-i, Supp.Fig 3c-e)**.

In the lipid category, fatty acid synthesis genes such as Acetyl-CoA carboxylases (Acaca, Acacb) and Elongases (Elovl6, Elovl2) were upregulated in 3xTg-AD mice on Control diet, while genes regulating fatty acid degradation such as Acetyl-CoA acetyltransferases (Acat1, Acat 3) and Acyl-CoA dehydrogenase medium chain (Acadm) that is crucial for MCFAs beta-oxidation were downregulated **(Fig 4c, 4i)**. MCT alleviated the aberrant expression of 40% (46/110) and CFHF of 57% (63/110) of these lipid-associated DEGs **(Supp.Fig 3c)**.

Our analysis further indicated that 3xTg-AD mice had downregulated expression of genes involved in ketogenesis and ketolysis (Bdh1, Hmgcs1, Hmgcs2, Hmgcl, Ehhadh, Hadh, Acat 3) and upregulated expression of numerous genes involved in carbohydrate-derivative processes, particularly in pyruvate metabolism (Adh1, Pklr, Aldh1b1) and pentose phosphate pathway (Pgd, Fbp1, G6pdx, Rpia)(**Fig 4g-h**). CFHF strongly upregulated ketone associated genes and downregulated 66% (28/42) of carbohydrate related genes while MCT had less pronounced effects (**Fig 4g-h, Supp.Fig 3c**).

Additionally, we identified AD-induced pathway changes for mitochondrial membranes and xenobiotic metabolizing processes. Mitochondrial membrane DEGs were all downregulated in 3xTg-AD mice on Control diet and partially restored by MCT and CFHF (**Supp.Fig 3d**). Xenobiotic metabolizing processes, including glutathione S-transferase gene family (Gstm1/2/3/6/7, Gstp3, Gstt3), UDP-glucuronosyltransferases (Ugt1a5/9, Ugt3a1, Ugt2b35) and ABC transporters (Abcb1b, Abcb1a, Abcc4-8) that are key elements in bile secretion and closely related to arachidonic acid and cholesterol metabolism, were deregulated in 3xTg -AD mice **(Supp.Fig 3e)**. More than 40% of DEGs associated with these processes were modulated by MCT or CFHF diets **(Supp.Fig 3c).**

Interestingly, closer analysis showed a particularly pronounced effect of CFHF diet on a subset of AD-induced DEGs whose expression was not just normalized but completely inversed versus WT mice (Fig 4c, 4e, **Supp.Fig 3f**). 26/844 genes that were either up- regulated or down-regulated in 3xTg-AD mice on Control diet, were respectively down- or up- regulated in 3xTg-AD mice on CFHF diet (Fig 4c, 4e, **Supp.Fig 3f**). Notably, this includes the Leptin Receptor gene (Lepr), which was up-regulated in 3xTg-AD mice on Control diet and decreased by CFHF to a lower level than WT mice on Control diet. Conversely, the gene encoding the fat storage inducing transmembrane protein 1 (Fitm1) was down-regulated in 3xTg-AD mice on Control diet and up-regulated by CFHF (Fig 4c, 4e **Supp.Fig 3f**).

These findings reveal the liver as a prominent site of AD-associated dysregulation and demonstrate that the dysregulated hepatic transcriptome is largely corrected by KI.

### MCT and CFHF drive distinct alterations on lipid and immune genes selectively in AD mice

In addition to partially correcting the liver’s AD-induced gene expression abnormalities, MCT and CFHF diets also elicited “new” DEGs. 414 DEGs were induced by the MCT diet in WT or 3xTg-AD mice versus the respective mice on the Control diet **(Fig 5a)**. This included 21 DEGs regulated in both WT and 3xTg-AD mice and that were enriched in GO pathways for transmembrane transport and lipid/cholesterol/sterol metabolism **(Fig 5b)**. 278 genes were altered by MCT uniquely in 3xTg-AD mice (**Fig 5a-c**) and GO analysis revealed that many of these genes were also involved in lipid metabolic processes, including downregulation of the fatty acid synthase (Fasn) gene that is essential for fatty acid synthesis, patatin-like phospholipases (Pnpla5, Pnpla3) known for regulating lipid droplets, and Hmgcr, the rate-limiting enzyme for cholesterol synthesis, and upregulation of Adipoq, which is associated with fatty liver (**Fig 5c, Supp. Table 5)**.

Even more prominently, 2039 new DEGs were induced by the CFHF diet in WT or 3xTg- AD mice (**Fig 5d**). Most of these DEGs were related to lipid/fatty acids and organic acids catabolism suggesting an increased Krebs cycle (**Fig 5e, Supp.Fig 3g**). Notably, both 3xTg- CFHF and WT-CFHF show an upregulation of Tnfaip8l1 which plays an anti-steatosis function (Supp. Table 6). Among the pathways modulated in WT mice and 3xTg-AD mice on CFHF diet, we noted that only 3xTg-AD mice had altered immune system process (**Fig 5e**). This pathway accounted for 166/1322 genes (13% of genes only altered in 3xTg-AD mice on CFHF) associated with inflammation and activation of immune system, such as IL1b, Ccl2, Cxcl2/14 and Nfkb1 (**Fig 5f, Supp.Table 6**).

Thus, MCT and CFHF diets exert distinct effects on the hepatic transcriptome of 3xTg- AD mice, with MCT specifically downregulating lipogenesis and cholesterol synthesis genes, and CFHF inducing activation of the immune system and major lipid metabolic changes.

### Plasmatic changes associated with AD-induced peripheral disturbances and the differential effects of KI

Inter-organ communication is largely mediated by circulating mediators. Since 3xTg- AD mice presented metabolic abnormalities and hepatic alterations in genes associated with glucose and lipid processes, and as many of these changes were modulated by KI, we evaluated AD- and KI-associated changes in the levels of plasma metabolic hormones and fatty acids.

Immunoassay of 8 plasmatic hormones (ghrelin, gastric inhibitory polypeptide (GIP), glucagon-like peptide 1 (GLP-1), insulin, leptin, plasminogen activator inhibitor (PAI-1), resistin and glucagon) revealed that 3xTg-AD mice had higher levels of ghrelin, glucagon and especially PAI-1, which was twofold higher compared to WT mice on Control diet (**Supp. Table 5**). The increase in these three hormones in 3xTg-AD mice is in line with their elevated caloric intake and hyperglycemia: Ghrelin is an orexigenic hormone that can stimulate secretion of glucagon, a pancreatic hormone whose levels regulate hepatic glucose, and PAI-1 is an antifibrinolytic protein whose levels are linked to hyperglycemia and hypertriglyceridemia. Dietary challenges did not affect these hormone levels, however both KI increased leptin levels in 3xTg-AD mice **(Supp. Table 5)**. Since leptin normally decreases body fat, the pronounced CFHF-induced increase in Leptin combined with increased body weight and white adipose tissue and downregulation of hepatic leptin receptor expression is suggestive of a CFHF-specific leptin resistance.

Lastly, we used gas chromatography with flame ionization detector (GC-FID) on plasma of treated mice to determine the concentrations (**Supp.Table 7**) and relative percentages (**Fig 6**, **Supp.Table 6**) of 21 long-chain fatty acids. Total amount of plasma fatty acids was similar in mice on Control and MCT diets and increased 1.5-fold in mice fed with CFHF diet (**Supp.Table 7**). The ratio of omega-6:omega-3 fatty acids, a measure of pro-inflammatory/anti-inflammatory polyunsaturated fatty acids (PUFAs)^37,38^, increased from around 7:1 in the Control and MCT diet-fed mice to 9.5:1 in the CFHF-fed mice (**Supp.Table 6**). These changes in total amount of fatty acids and omega-6:omega-3 fatty acids reflect the composition of the administered diets (Supp. Table 6**, Supp.Table 2**) and indicate the direct impact of dietary fat levels on circulating fatty acids. By using relative percentages to compare specific acids, we observed that the genetic risk for AD associated with 3xTg-AD mice correlated with a subset of plasma fatty acid changes: increased C20:0, C18:1Ω9, C20:1Ω9 and decreased C18:2Ω6 and C20:5Ω3 (**Supp.Table 6 and Fig 6**). Arachidic acid (C20:0), the only saturated fatty acid that was changed, was higher in 3xTg-AD mice (**Supp.Table 6**). Oleic acid (C18:1Ω9), the main monounsaturated fatty acid (MUFA), and another omega-9 MUFA, eicosenoïc acid (C20:1Ω9), were higher in 3xTg-AD mice on the Control diet than WT mice (**Supp.Table 6**, **Fig 6f**), and the C18:1/C18:0 index, a measure of activity for the enzyme that converts saturated fatty acids into MUFAs, stearoyl-coenzymeA desaturase (Scd), was likewise higher in 3xTg-AD mice on the Control diet (**Fig 6d**) . 3xTg-AD mice on the Control diet also had a higher percentage of total MUFAs, a decreased percentage of total polyunsaturated fatty acids (PUFA), and an increased MUFA/PUFA ratio (**Fig 6a, Supp.Table 6**). Among the PUFAs, 3xTg-AD mice on Control diet presented lower levels of linoleic acid (C18:2Ω6) compared to WT mice. Interestingly, many of these differences between WT and 3xTg-AD mice on the Control diet were no longer significant when the WT mice on the Control diet were compared with 3xTg-AD mice on MCT or CFHF (**Fig 6a-b,d,f,g**). A subset of plasma fatty acids including omega-7 fatty acids and several PUFAs, was changed in opposing directions in MCT and CFHF fed mice when compared to Control diet. Cis-palmitoleic acid (C16:1Ω7) and vaccenic acid (C18:1Ω7) were both elevated by MCT and markedly reduced by CFHF (**Fig 6e**, **Supp.Table 6**). Among the omega- 6 fatty acids, dihome-γ-linoleic acid (DGLA, C20:3Ω6) was reduced by CFHF in both WT and 3xTg-AD mice (**Fig 5h**, **Supp.Table 3**) while docosapentaenoic acid (DPA, C22:5Ω6) levels were increased fourfold by CFHF compared to Control diet selectively in 3xTg-AD mice (**Fig 6i, Supp.Table 6**). The omega-3 fatty acid docosahexaenoic acid (DHA, C22:6Ω3) was reduced by CFHF in both WT and 3xTg-AD mice **(Fig 6k, Supp.Table 6)**.

**Figure 6:**
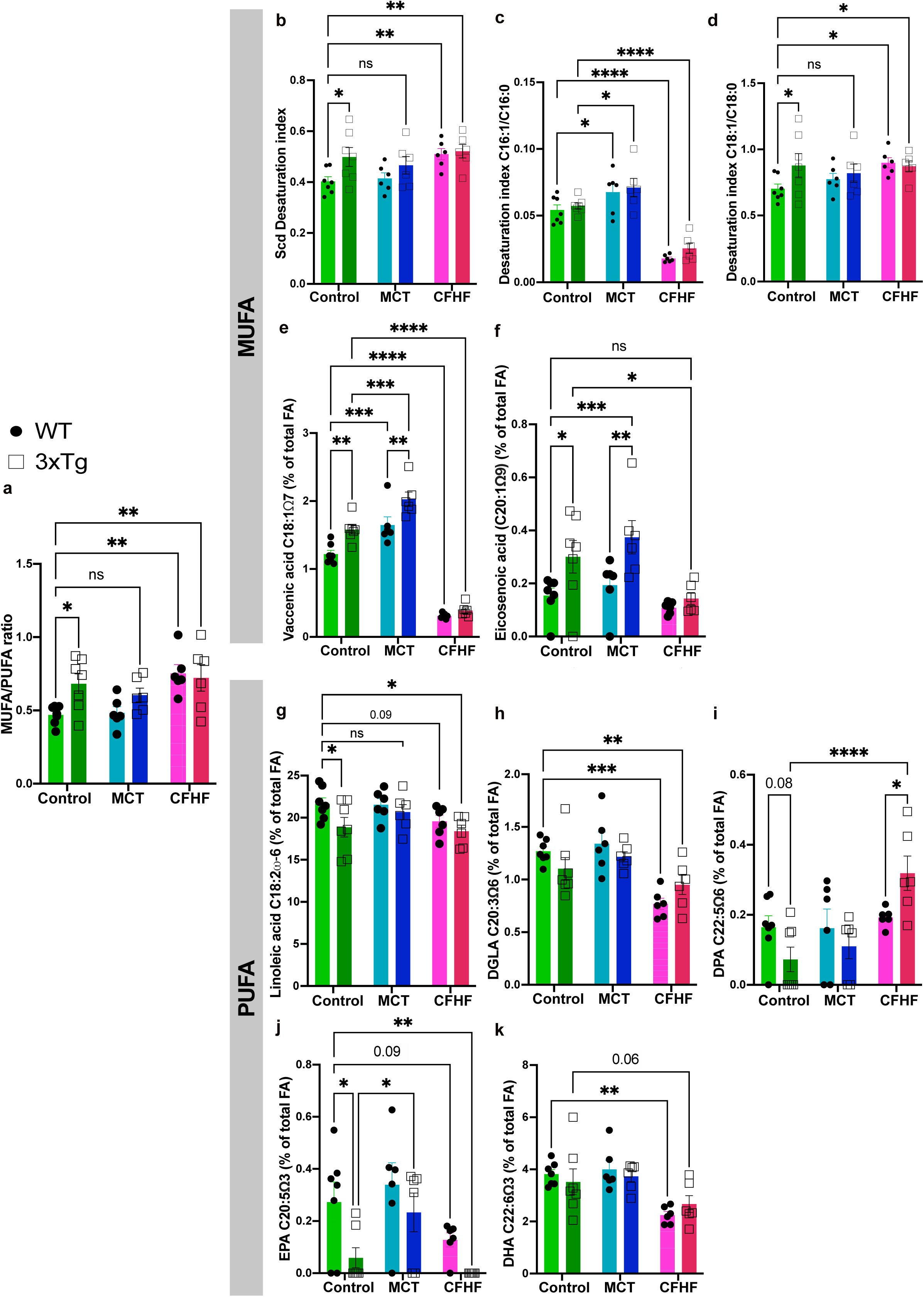
MCT and CFHF drive opposing plasma concentrations of omega-7 MUFAs and some PUFAs in AD mice. Plasmatic fatty acid levels were evaluated using gas-chromatography with flame ionization detector (GC-FID n=6-7/strain/diet). a) Ratio of mono-unsaturated and Poly-unsaturated fatty acids (MUFA/PUFA, 2-way ANOVA showing diet(p-value=0.0055, F(2,32)=6.142) and strain(p-value=0.0411, F(1,32)=4.531) effects. Fisher post hoc test (detailed on graph)) b) Total desaturation index due to Stearoyl-Coenzyme A 9-desaturase (Scd, 2-way ANOVA showing diet(p-value=0.0258, F(2,32)=4.110) and strain(p-value=0.0273, F(1,32)=5.347) effects) calculated with the desaturation index of (c)C16:0/C16:1 (2-way ANOVA showing diet(p-value=0.0258, F(2,32)=4.110) and strain(p-value=0.0273, F(1,32)=5.347) effects) and (d)C18:0/C18:1 (2-way ANOVA non-significant). Fisher post hoc test (detailed on graph)) e-f)MUFAs significantly affected by diet and/or strain : e)Vaccenic acid (2-way ANOVA showing diet(p-value<0.0001, F(2,32)=198) and strain(p-value<0.0001, F(1,32)=19.87) effects). f)Eicosenoic acid(2-way ANOVA showing diet(p-value=0.0042, F(2,32)=6.528) and strain effects (p-value=0.0018, F(1,32)=11.54)), Fisher post hoc test (detailed on graph)) g-i)Omega-6 PUFAs significantly affected by diet and/or strain: g)Linoleic acid (2-way ANOVA showing nearly significant diet effect (p-value=0.0657, F(2,32)=2.969) and strain effect(p-value=0.0283, F(1,32)=5.277)). h)DGLA (2-way ANOVA showing diet effect(p-value<0.0001, F(2,32)=14.33) with trend toward interaction between strain and diet (p-value=0.0817, F(2,32)=2.711)). i)DPA (2-way ANOVA showing significant diet effect(p-value=0.0022, F(2,32)=7.442) with interaction between strain and diet (p-value=0.0186, F(2,32)=4.522)) j-k)Omega-3 PUFAs significantly affected by diet and/or strain: j)EPA (2-way ANOVA showing significant diet(p-value=0.0037, F(2,32)=6.718) and strain effects(p-value=0.0042, F(1,32)=9.518)), k)DHA (2-way ANOVA showing diet effects (p-value=0.0002, F(2,32)=11.52)) Fisher post hoc test: *p-value<0.05; **p-value<0.01; ***p-value<0.001; ****p-value<0.0001

These plasmatic data identify biomarkers and potential mediators of AD-induced peripheral metabolic alterations and the differential effects of MCT and CFHF.

## Discussion

Using two well-known models of AD, this study provides several insights into the central and peripheral targets of KI in AD.

### KI improve hippocampal function and dendritic structure of AD mice

Both KI that we used improved the hippocampal-dependent learning and memory performance of AD mice, and this was associated with increased density of dendritic spines within the dentate gyrus. These functional and structural improvements were generalizable to both the 3xTg-AD and 5xFAD models. This demonstrates reversal of cognitive deficits induced by amyloid-beta overexpression by KI and identifies a potential central basis for this effect. The results corroborate clinical reports^23,30,39–41^ and are in line with evidence that ketogenic diet improves long-term potentiation in AD mice ^42^. The absence of diet effects on CA1 neurons in this study highlights that specific hippocampal neuronal populations are likely to be particularly sensitive to KI.

Notably, the central effects of the MCT diet on the hippocampus do not appear to hinge on achieving a significant or sustained elevation of circulating ketones. The MCFAs in MCT can potentially directly affect the brain independently of ketones, as they can cross the blood brain barrier^29,43^ and themselves constitute bioenergetic substrates^22^. They can also improve mitochondrial biogenesis and modulate glutamate excitotoxicity ^22,44^, which can contribute to the preservation of synaptic connections. Similar to our observations, Ota et al,^39^ reported no increase in circulating ketone bodies in patients receiving MCT treatment during a long-term intervention, whereas an acute intervention in the same subjects triggers ketosis. MCT administration induces a transient increase of ketone bodies cleared from the blood within 2-3 hours^25,45,46^. Therefore, long-term “mild intermittent ketosis”^47^ might also play a role in mediating the cognitive effects of the MCT diet.

### AD genetic risk triggers peripheral metabolic disturbances linked to hepatic alterations

The 3xTg-AD model exhibits systemic metabolic abnormalities prior to the formation of plaques and neurofibrillary tangles^48,49^, including visceral fat accumulation, weight gain and impaired glucose tolerance^49,50^. Importantly, here we identify the liver as a key target and potentially mediator of AD-induced peripheral defects. The hepatosplenomegaly of 3xTg- AD mice was associated with high levels of hepatic lipid droplets, indicating hepatic steatosis, as well as substantial transcriptomic deregulation that included genes involved in fatty acid, glutathione, bile acids, carbohydrate and mitochondrial pathways. Transcriptomic analysis of 5xFAD mice liver also revealed alterations in lipid metabolism especially cholesterol (data not shown). The enlargement of the spleen is regarded as an indicator of liver disease and increased inflammation and has been observed in both 3xTg- AD and 5xFAD models of AD^49,51–54^. Interestingly, non-Alcoholic Fatty Liver Disease (NAFLD) is thought to increase the risk to develop dementia^55–57^. Our data suggest that dementia risk factors (APP, PSEN1, Tau mutations) can induce minor steatosis in the liver and alter hepatic processes. In line with this, changes consistent with altered hepatic functions have been observed in AD patients^14,16^ such as altered plasma levels of liver enzymes^13,16^ and bile acids^12,58^. Altered brain and blood glutathione levels were also observed in AD patients^59^. We observed that genes involved in pyruvate formation and pentose phosphate pathways were upregulated and Insulin receptor substrate 2 (Irs2) was downregulated in 3xTg-AD mice, suggesting alterations in hepatic glycolysis or gluconeogenesis and in response to insulin. Also downregulated were lactate dehydrogenase (Ldha) and pyruvate carboxylase (Pcx), which catalyze lactate/pyruvate conversion and oxaloacetate synthesis from pyruvate ^60^, the latter being involved in anaplerotic pathways notably gluconeogenesis under carbohydrate- restrictive conditions^61,62^. These changes are in line with the impaired peripheral glucose response and increased glucagon levels observed in our mice.

Interestingly, among the deregulated hepatic genes deregulated were numerous genes previously identified in GWAS studies as AD risk genes such as Chrna2, Psen1, Bin1, Ercc2, Ptk2b, Cox7c, Tpcn1 and Klf16^63–65^. Two-pore channels such as Tpcn1 and the transcription factor Klf16 were also associated with NAFLD^66–68^, suggesting an interaction among genetic risk factors of AD in a potential liver-brain axis. Many of these AD-genes were restored by the KI (Bin1, Ercc2, Tpcn1 and Klf16).

### Complex interplay between AD genetic risk and ketogenic interventions

This study provides substantial data that in addition to its neuroprotective changes in the brain, MCT supplementation has effects on food intake, body fat, improving glucose metabolism, regulating plasma fatty acid profile and hepatic transcriptome. The CFHF diet showed similar effects on the brain compared to MCT yet modified the peripheral phenotype of 3xTg-AD mice in largely opposing directions. 3xTg-AD mice and their WT counterparts also showed differences in their responses to CFHF, with 3xTg-AD mice showing hyperleptinemia, reduced ketosis, reduced diet-derived anti-steatosis effect and upregulated pro-inflammatory genes in the liver. Thus, challenge with the CFHF diet revealed a metabolic vulnerability in 3xTg-AD mice consistent with an altered ability to metabolise a high proportion of long-chain triglycerides.

Monitoring peripheral metabolism is highly relevant in AD. Glycemia and insulin sensitivity can predict the risk of dementia or cognitive decline^69,70^ and treatments regulating metabolic defects such as antidiabetic drugs showed potential to reduce dementia risk^71^. In line with our observations, MCT diet reduces white fat^72^ in obese patients and prevents glucose dysregulation in individuals subjected to high-fat diet^73^. The faster hepatic oxidation of MCT, compared to long-chain triglycerides (LCT), is thought to prevent lipid uptake in adipose tissue^72^. Mouse studies show that MCT improves insulin sensitivity by regulating insulin-modulating hormones such as glucagon-like peptide 1 (GLP-1) and gastric inhibitory polypeptide^74,75^. Although, these hormones were not altered in response to MCT in this study, increased hepatic expression of Irs2 and decreased Hmgcr were observed. This suggests potential effects on hepatic glucose metabolism and cholesterol synthesis, two pathways closely related^76^ which affect peripheral glucose response. Reduced liver cholesterol in animals fed with MCT-based interventions have also been reported^77,78^.

While KI are often associated with weight loss, CFHF-induced fat increases and hyperglycemia have been reported in healthy human and non-transgenic mouse studies. These effects are either attributed to physiological adaptation to the diet which should be temporary^79^ or to the very-low proportion of carbohydrates^80^ which is considered detrimental for long-term. An increased metabolic vulnerability similar to what we report in 3xTg-AD mice has been described for brain glucose metabolism in familial AD patients, but a similar observation in the periphery remains a hypothesis^81,82^.

### Novel mediators/effectors of systemic changes induced by AD and KI

Our study opens the door to investigating the roles of several molecules as potential mediators and/or effectors of the systemic changes induced by AD genes and/or KI.

The metabolic hormones ghrelin, glucagon, and PAI-1 were increased in the plasma of 3xTg-AD mice, alongside with disturbed plasma fatty acid profile and alterations in the liver. Interestingly, the metabolic abnormalities in 3xTg-AD mice resemble features of T2D and obesity, reinforcing the idea of a pathophysiological link between AD and peripheral systemic abnormalities^83,84^. Our data suggest that these abnormalities can derive from familial AD mutations. Although a causal link between familial AD and systemic disabilities is not clinically established, studies suggest that it can involve amyloid beta^81,83,85,86^. For instance, the peripheral overexpression of APP and the injection of amyloid beta oligomers which precede senile plaques in the brain are both sufficient to produce peripheral metabolic defects related to glucose and lipid processes^82,85,87,88^. Increased levels of PAI-1 were also associated with brain amyloid deposits in healthy individuals with high risk to develop dementia^86,89^

The MUFA/PUFA ratio was significantly increased in 3xTg-AD mice due to increased oleic acid (C18:1Ω9) and decreased linoleic acid (C18:2Ω6) which are the most abundant fatty acids in their categories, as well as lower levels of EPA compared to WT mice. This parallels the increase of MUFA-synthetic enzymes in the liver (Scd1, Elovl6). C18:1Ω9 was associated with AD brain lipid accumulation^90^ and the inhibition of its production from stearic acid with an inhibitor of Scd shows therapeutic potential for AD and Parkinson’s disease^34,91,92^. C18:1 Ω9-induced brain lipid accumulation was firstly described in 3xTg-AD mice and proved to impair neurogenesis^90^. The observed increase in its plasma levels might lead to systemic alterations. Concerning PUFA, lower levels of omega-3 fatty acids such as EPA have also been reported in AD patients^93,94^.

MCT increased levels of omega-7 MUFAs such as palmitoleic acid (C16:1Ω7) and vaccenic acid (C18:1Ω7) while CFHF strongly reduced them compared to Control diet. C16:1Ω7 is synthesized by Scd^92^ from C16:0 and C16:1Ω7 can be converted into C18:1Ω7 by the elongation of very long-chain fatty acid member 6 (Elovl6) and Elovl5 which also elongate C18 PUFAs^95–97^. Interestingly, the hepatic expression of these enzymes, excepted Elovl5, are reduced by both KI. Low peripheral serum levels of C16:1Ω7 were observed in AD patients^98^ and C16:1Ω7 has potential benefits for obesity and T2D^99,100^, while C18:1Ω7 is associated with lower risk of diabetes^101^. Interestingly obesity triggered by a high-fat diet incorporating coconut oil results in elevated C18:1Ω7 levels, yet without inducing hyperglycemia or insulin resistance ^102^, indicating that alterations in the fatty acid profile induced by an MCT source may mitigate the metabolic disturbances associated with obesity, which is in line with our observations.

Leptin signaling and Scd activity also appear as mediator of KI, especially in low-carbohydrate/high-fat diet. CFHF induced a strong reduction of the hepatic expression of Scd1 and Lepr. It also reduced hepatic lipid droplets indicating anti-steatosis effects, previously reported in human^103^ and animal studies^104^. Brain leptin reduces hepatic steatosis and hepatic Scd1 activity^105,106^. Another study supports that low-carbohydrate/high fat diet effects on body weight requires increased leptin levels and functional leptin signaling^107^.

### Limits & Strengths

This study has certain limitations. We did not monitor the acute rise in plasma ketones following MCT ingestion, which was provided ad libitum rather than as single boluses. This would have allowed the evaluation of any transient increase of ketones after MCT intake. Second, we did not control for the metabolic state while collecting plasma for the analysis of metabolic markers. Metabolic markers are sensitive to food intake and a fasting of 1 or 2h would have led to less variability and eliminated the possibility of postprandial bias. Third, we did not investigate the incorporation of circulating fatty acids into the downstream classes of complex lipids they contribute to. In future studies this will provide a more complete understanding of the plasma lipidome allowing for example the distinction of fatty acids within esterified cholesterol versus the ones in triglycerides. Nonetheless, this study offered an overview of the central and peripheral effects of KI in AD mice. It employed various techniques to analyze the peripheral metabolism with enough animals. Administrating the dietary interventions in non-AD mice highlighted the effects of such interventions in physiological conditions and revealed latent metabolic deficits in AD mice. The use of two models permitted the analysis of the diet response at different stages of AD pathophysiology.

Ketogenic interventions are emerging as promising strategies for AD and other neurodegenerative conditions^18^. The mechanistic insights into central and peripheral targets of KI provided here will help contribute to the rational design of more optimized and robust therapeutic strategies.

## MATERIALS AND METHODS

### Mice and treatment

#### Ethics

All experiments were carried out following the guidelines of the Canadian Council of Animal Care and were approved by the institutional animal care committee of the University of Montreal Hospital Research Center (CRCHUM) and the University of Sherbrooke.

#### 5xFAD mice

The generation and characterization of the model were previously described ^32,108^. The 5xFAD (Jackson laboratory RRID:MMRRC_034840-JAX) transgenic mouse model carries five familial Alzheimer’s disease (FAD) mutations, including **Swedish (K670N, M671L), Florida (I716V), and London (V717I)** mutations in the APP transgene, as well as **M146L and L286V FAD** mutations in the PSEN1 transgene, both under mouse Thy1 promoter. These transgenic mice were bred onto the B6SJLF1/J background (Jackson laboratory stock#: 100012). For our experiments in this study, we utilized both heterozygous mutant mice and non-carrier mice as appropriate controls.

#### 3xTg-AD mice and B6;129 strain wild type

The generation and characterization of the model were previously described^33,109^. 3xTg-AD mice (Jackson laboratory, RRID:MMRRC_034830-JAX) possess three human dementia- causing mutations that lead to AD’s pathological traits: **APP_Swe_ and tau_P301L_** encoded in two independent transgene constructs under control of the central nervous system (CNS) mouse regulatory element Thy1.2, co-microinjected into single-cell embryos harvested from homozygous mutant **PS1_M146V_** knock in (PS1-KI) mice. Wild type mice are B6;129 (WT, the 3xTg-AD background strain, Jackson laboratory stock#: 101045) obtained from a cross between C57BL/6J females (B6) and 129S1/SvImJ males (129S).

#### Housing and Dietary interventions

All included mice were females, organized in three cohorts, each with specific treatment timelines:

First Cohort: 5xFAD and control mice received a one-month dietary intervention from 6 to 7 months of age, evaluating cognitive recovery.

Second Cohort: 5xFAD mice were on diets for three months, spanning ages 2 to 5 months, 3xTg-AD mice were on diets for five months, spanning ages 3 to 8 months to analyze protective changes at pre-symptomatic level.

Third Cohort: 3xTg-AD and WT mice were fed diets for six months starting at 3 months of age to assess peripheral metabolic changes.

Mice used for cognitive and brain analysis were group-housed (3 to 4 per cage). The one used for metabolic analysis were single housed for food intake monitoring. All animals were in reverse cycle room (12h Dark-light cycle 10AM – OFF, 10PM – ON) with food/water ad libitum. Mice received either a Control diet (C, 70% carbohydrate, 20% long-chain triglycerides, 10% protein, 4.1 kcal/g, Research Diets product #D17121209I), a Control diet enriched with medium-chain triglycerides (MCT, 70% carbohydrate, 10% long-chain triglycerides, 10% medium-chain triglycerides (caprylic acid to capric acid ratio of 3:2), 10% protein, 4.1 kcal/g, Research Diets product #D17121210I) or a carbohydrate-free high fat diet (CFHF, 90% long-chain triglycerides, 10% protein, 6.7 kcal/g, Research Diets product# : D10070801).

#### Morris water maze

The water maze comprises a 100 cm diameter pool surrounded by additional cues defining four quadrants (NE, NW, SE, SW). The water temperature was maintained at 23 ± 1 °C. The tracking software used was EthoVision XT 17. Mice underwent training with 4 trials per day over 4 days. On Day 5, a probe trial was conducted with the platform removed. Starting positions were determined using a pseudorandom algorithm and varied for each trial. The maximum trial duration was 60 s, and the software recorded mice as having reached the platform if they spent 2 s in the platform zone. Mice could remain on the platform for 20 s, and if they didn’t find it, they were gently guided there and allowed to stay for 20 s. After each trial, mice were lightly dried and placed on a heating pad until the next trial. The inter-trial interval ranged from 15 to 30 min.

#### Golgi Cox staining

The experiment and analysis were performed as previously described^34^. Mice were deeply anesthetized with isoflurane and promptly decapitated. The brain was extracted, and one hemisphere was sliced 5 mm from the olfactory bulbs and 4 mm from the brain stem using a mouse brain mold, exposing the hippocampus. Golgi staining was performed using the Slice Golgi Kit (Bioenno Tech, USA). Slices between Bregma −1.06 to −2.20 mm were post- fixed initially in fixative solution for 1 h at 4 °C, followed by an additional 48 h at 4 °C in fresh fixative solution before slicing on a vibratome at 150 µm. The sections were washed three times with PB pH 7.4. Free-floating sections were subsequently stained in the impregnation solution for 24 h and then in a fresh impregnation solution for an additional 4 days at room temperature on a rocker. Next, the sections were washed once with PBS-T to remove the impregnation solution, followed by further washes with PBS 1× pH 7.4. Golgi staining was performed by immersing sections in solution C for 1 min and then in post-staining solution D for 2 min. Sections underwent dehydration in a series of ethanol baths (50%, 80%, 95%, and 100%), followed by two Xylene washes. Finally, the sections were mounted with Permount (Thermo Fisher Scientific, Canada) and allowed to air-dry overnight before imaging.

Dendritic spine density was quantified by counting spines present on secondary and tertiary dendrites of neurons in the dentate gyrus and the basal dendrites of neurons in the CA1 region of the hippocampus. This assessment was conducted using a light microscope at ×100 equipped with a color camera (Olympus BX43).

Quantifications in DG or CA1 in dorsal hippocampus (Bregma -1.06mm to -2.70mm) included approximately two neurons per region (6-12 sections per animal n = 5-6 mice per treatment group) for a total of ∼50-90 neurons per group. All quantifications were done by a blinded experimenter.

### Measures of systemic metabolic parameters

#### Daily caloric intake and body weight gain per caloric intake

The food pellets were changed twice a week according to the food company recommendations. Food intake(g) was calculated by subtracting left food in the cage from provided food. Then this amount of consumed food was multiplied by the energy weight of each diet and divided by the number of days before a food change to obtain the daily caloric intake (kcal). The mice were weighed once or twice a month throughout the experiment. The average daily body weight gain was calculated as the difference of two measures divided the number of days between these measures. The body weight gain per caloric intake (g/kcal) is the ratio of daily body weight gain out of daily caloric intake.

#### Blood glucose and beta-hydroxybutyrate monitoring

At different time points, circulating glucose and beta-hydroxybutyrate (BHB) levels were measured using the freestyle lite blood glucose monitoring system (Abbott, product#: ART20771) with the corresponding reading strips (Abbott, REF#: 70825-70) and the freestyle precision neo (Abbott, product# : ART30531)with the blood BHB test strips (Abbott, REF# : 70748-75). The mice were pricked in the saphenous vein with a needle to have the required blood drops for the strips.

#### Body composition analysis

##### -Micro-CT

The micro-computed tomography scan (*In vivo* Micro-CT SkyScan 1176 microtomograph, Bruker) was used to visualize subcutaneous and abdominal peripheral fat distribution in live animals under anesthesia (induced by isofurane 3% and Oxygen 1-1.5 L/ml and maintained by isofurane 3% and Oxygen 0.75-1L/ml). The animals were placed back down on the animal bed and the Micro-CT was set according to company requirements (Quantifying adipose tissue (fat) in a mouse or rat by in-vivo microCT, Bruker microCT). The values set for performing Micro-CT were as follow : X-ray applied voltage : 30kV / position of sample bed: - 79.431mm / spatial resolution: 4000 X 2672 pixels or 9mm pixel size / X-ray filter: AI 1mm. The volume of interest was the whole mouse body. It was obtained by the sum of region of interests delineated on 2D images. On 2D images, the mouse body boundary was selected with a binary threshold of 22-255 greyscale. Other binary thresholds were applied to delineate fat (13-21 greyscale) and bones (90-255 greyscale).

##### -EchoMRI

Total fat mass, lean mass and total water mass in live mice unanesthetized, were measured using nuclear magnetic resonance (EchoMRI^TM^ – Whole body composition analyzer). The scan lasts 3 minutes/mouse and was performed on days 0, 28, 56, 112 and 168 during the diet experiment.

#### Glucose / Insulin Tolerance Test

##### -Glucose Tolerance test

After 16 hours fasting to normalize blood glucose level, we performed an intra-peritoneal (IP) injection of dextrose 50% (concentration injected: 1g/kg animal body weight, DIN: 02468514, Pfizer product#: 00409-6648-16) and measured blood glucose evolution each 15 minutes for 1h30 minutes (as described above for *Blood glucose monitoring*)

##### -Insulin Tolerance Test

After 1 hour fasting to normalize blood glucose level, we performed an intra-peritoneal (IP) injection of Humulin R (human insulin, concentration injected: 0.5U/kg animal body weight, DIN: 00586714, Eli Lilly) and monitored blood glucose evolution each 15 minutes for 1h30 minutes.

#### Comprehensive Lab Animal Monitoring System (CLAMS)

The cages are placed in an environment where light and temperature are controlled - 12 hours of light, 12 hours in the dark at 21 °C. The energy expenditure, gas exchanges and ambulatory activity of the animals were measured using the CLAMS system (Columbus Instrument, over a period of 48 hours (24 hours of habituation, 24 hours of measurement). The software CLAX is used to extract the data.

### Blood sampling and plasma collection

Following isoflurane anesthesia, blood was collected from the inferior vena cava (IVC). An incision along the abdominal region exposed to the IVC beneath the intestines. Blood from the IVC was slowly drawn using a 27-gauge needle attached to a syringe, maintaining a horizontal needle orientation to prevent vein damage. The blood was swiftly transferred to EDTA tubes to prevent coagulation. After mixing by inverting and flicking the tubes, blood from the EDTA tubes were centrifuged at 1000g for 20 minutes at 4°C to separate plasma, which was carefully collected into new Eppendorf tubes. The plasma samples were temporarily stored on ice and then stored at -80°C for subsequent analysis.

### Gas-chromatography with flame ionization detector (GC-FID)

Total lipids were extracted from plasma samples using the Folch method^110^. Sample weights (2.31–5.31 mg) were used to calculate the amount of standard (30–70 µg) of 1-O- Heptadecyl-2-Acetyl-sn-Glycero-3-Phosphocholine (17:0 PC; Sigma-Aldrich cat# 850360P) previously diluted in 10 mL of chloroform was added as an internal standard prior to extraction. Saponification of the fatty acids was performed using KOH-methanol, protonation followed with 12 N HCl and methylation with 12% BF_3_-methanol (Thermo Fisher Scientific cat# AC402765000), as described previously in Plourde et al.^111,112^

The fatty acid composition of the plasma samples was analyzed by gas chromatography coupled with a flame ionization detector (model 6890; Agilent). 1 µl of the sample was injected in splitless mode at 250 °C. The methylated fatty acids were carried by helium at a pressure of 107 kPA through a 50 m BPX-70 fused capillary column (0.22-mm diameter, 0.25-µm film thickness; SGE). A thermal gradient was applied on the column as described in Chevalier et al.^113^. Fatty acids were then detected at the end of the column by the flame ionization detector at 260 °C. The total run time was 61.75 min per sample. Nu-Check-Prep standards (GLC-569) were used to identify 45 different fatty acid methyl esters and the chromatogram analysis was performed using OpenLab CDS ChemStation version C01.10.

### Immunoassay of metabolic markers

Plasma samples was used to perform the Bio-Plex Pro Mouse Diabetes 8-plex assay (Biorad #171F7001M) following the manufacturer’s protocol. This multiplex assay can detect the following molecules: ghrelin, gastric inhibitory polypeptide (GIP), glucagon-like peptide 1 (GLP-1), insulin, leptin, plasminogen activator inhibitor (PAI-1), resistin and glucagon. The instrument used for detection was Bio-Plex MAGPIX system.

### Histological analysis of the liver

#### Oil redo O (ORO) and Boron-dipyrromethene (BODIPY)

Mice were anesthetized by isoflurane and decapitated. The largest liver lobe was collected and post-fixed in 4% paraformaldehyde for 24hours. Transverse sections of 40 μm thickness were obtained from liver tissue using a vibratome (Leica Model VT1000S). The sections were stored in antigel (40% Phosphate Buffer Saline (PBS), 30% Glycerol, 30% Ethylene Glycol) at -20°C until analysis. *<u>ORO:</u>* The tissue sections were washed, first with PBS, and then 60% isopropanol. The sections were stained with ORO (0.005%ORO (w/v) in isopropanol) diluted in distilled water with a factor of 0.6. They were rinsed again with 60% isopropanol, distilled water and mounted onto microscope slides using MOWIOL (13% (w/v) polyvinyl alcohol and 2% (w/v) DABCO in 2:1 Tris-Base (pH 8.5): Glycerol). Qualitative analysis of lipid droplet sections was performed blindly using a light microscope equipped with a color camera (Olympus BX43). *<u>BODIPY:</u>* The tissue sections were washed, first with PBS, and stained with a solution of BODIPY and Hoechst diluted in PBS (Hoechst 1:500, BODIPY 1:1000). The sections were mounted on microscope slides using MOWIOL and dried overnight protected from light. Qualitative analysis of lipid droplet sections was performed blindly using a fluorescent microscope.

#### Hematoxylin and Eosin (HE) staining

For HE staining, tissues were deparaffinized in xylene, rehydrated, and incubated in hematoxylin for 2 min, rinsed in water, dipped quickly in 0.5% acid, and washed in water. Tissues were then immersed in 0.2% NaHCO3, rinsed in water, dipped in eosin for 2 min and briefly rinsed in water before dehydration and mounting. Qualitative analysis of lipid droplet sections was performed blindly using a light microscope equipped with a color camera (Olympus BX43). Lipid droplet levels were quantified using ImageJ and protocol mentioned here ^114^. Initially, the images were converted into 8-bit grayscale. Then, a black-white inversion was applied, followed by adjustments to the detection threshold that emphasize the droplets. Finally, particle analysis was utilized to select and quantify the lipid droplets(size>20 pixel^2^, circularity: 0.75-1.00).

### Whole Liver RNA sequencing

#### RNA extraction, Library prep and Sequencing

RNA-extraction was performed as previously described^34^. Mice were anesthetized with isoflurane and decapitated. Piece of liver was dissected, immediately frozen on dry ice and conserved at -80°C until experimentation. Total RNA was extracted with TRIzol (Invitrogen) and chloroform (Invitrogen). RNA was then purified with the RNeasy mini kit (Qiagen) according to manufacturer’s instructions. 500ng of total RNA was used for RNA-sequencing libraries preparation. RNA quality control was assessed with the Bioanalyzer RNA 6000 Nano assay on the 2100 Bioanalyzer system (Agilient technologies) and all samples had a RIN above 8. Library preparation was done with the KAPA mRNAseq Hyperprep kit (KAPA, Cat no. KK8581). Ligation was made with 38 nM final concentration of Illumina dual-index UMI (IDT) and 11 PCR cycles was required to amplify cDNA libraries. Libraries were quantified by QuBit and BioAnalyzer DNA1000. All libraries were diluted to 10 nM and normalized by qPCR using the KAPA library quantification kit (KAPA; Cat no. KK4973). Libraries were pooled to equimolar concentration. Sequencing was performed with the Illumina Nextseq500 using the Nextseq High Output 75 cycles kit using 2.6 pM of the pooled libraries. Around 16-27 M single-end PF reads were generated per sample. Library preparation and sequencing was made at the Institute for Research in Immunology and Cancer’s Genomics Platform (IRIC).

#### Bioinformatics

RNA-sequencing analysis was performed using bioinformatic pipelines available in Dr Tetreault’s laboratory and run on clusters from the Digital Research Alliance of Canada (computational resources assigned to Dr Tetreault’s team). Sequences were trimmed for sequencing adapters and low quality 3’ bases using Trimmomatic version 0.35^115^ and aligned to the reference mouse genome version GRCm38 (gene annotation from Gencode version M23, based on Ensembl 98) using STAR version 2.5.1b^116^. Differential expression analyses were performed with the Bioconductor R package TCC v1.24.0^117^. Group data was analyzed pairwise and normalized using the DEGES/DESeq2 generalized linear model. Differentially expressed genes were identified with the TCC implementation of likelihood ratio test within DESeq2 with a p-value < 0.01 threshold. Gene set analyses were accomplished on called DEGs using ShinyGO v0.741: Gene Ontology Enrichment Analysis for biological process, molecular function and KEGG^118^.

Volcano plots were generated with GraphPad Prism 10.0, barplots were generated with R package ggplot2 (H. Wickham. ggplot2: Elegant Graphics for Data Analysis. Springer-Verlag New York, 2016.)

Heatmaps generated with R package Pretty Heatmaps (Raivo Kolde (2019). pheatmap: Pretty Heatmaps. R package version 1.0.12. https://CRAN.R-project.org/package=pheatmap). Venn diagramms were used to compare contrasts of interest and generated with Deep Venn (Tim Hulsen (2022). <u>arXiv:2210.04597</u>).

### Gene categories

For identification of the gene categories: “Lipid”, “Ketone”, “Carbohydrate”, “Mitochondrion” or “Xenobiotic”, gene set analysis was first performed on DEGs using ShinyGOv0.741 and Gprofiler. Then, gene categories relating to Lipids (lipid, fatty acid, steroid, linoleic acid, arachidonic acid, PPAR signaling), Ketone (Butanoate, ketone bodies), Carbohydrate (pyruvate, carbohydrate, pentose phosphate), Mitochondrion(mitochondrion, mitochondrial), Xenobiotic(Bile, drug, xenobiotic, ABC transporters, glutathione, glucuronidation) were formed using the keywords in parentheses.

### Statistical analysis

Statistical analyses were performed using GraphPad Prism 8.0 and 10.0(GraphPad Software). Unpaired t-test and two-way ANOVA with Uncorrected Fisher’s LSD test were used for appropriate situations. Data are presented as means ± standard errors of the mean. Statistical tests performed are detailed in the legend for each figure. P-values less than 0.05 were considered statistically significant and p-values between 0.05 and 0.1 were considered nearly significant.

## Author contributions

P.E.H.M organized the team, carried out the experiments, analyzed data, and co-wrote the manuscript. L.K.H, C.L.M, F.P., A.A. and M.M. performed dissections and helped to monitor weekly measures of food intake and body weight. L.K.H, S.M., M.G. helped to perform IPGTT and IPITT. G.M.B and M.T. performed the bioinformatic analysis. E.B helped to perform the immunoassay. M.B. helped to obtain brain slices for Golgi staining. A.C. analyzed liver histology. M.A., A.V. and M.P. performed GC-FID. M.T. contributed to review the manuscript. K.J.L.F developed the concept, analyzed data, and co-wrote the manuscript. All authors read and approved the final manuscript.

## Acknowledgement

We thank the Genomics Platform of the Institute for Research in Immunology and Cancer (IRIC) for the bulk liver sequencing, Electron Microscopy and Histology Research Core of the Faculté de Médecine et des Sciences de la Santé at the Université de Sherbrooke for the Hematoxylin Eosin staining and the paraffin liver slices, Compute Canada (www.computecanada.ca), GenAP (genap.ca) and its private Galaxy instance (galaxy.org) for bioinformatic resources, Pr Kaufmann lab for their MAGPIX system and the platform of small animal phenotyping and imaging of CHUM research center for the EchoMRI, Micro-CT scan and calorimetric cages . This work was funded by operating grants to K.J.L.F. from the Canadian Institutes of Health Research (CIHR), the Alzheimer Society of Canada, and the Natural Sciences and Engineering Research Council (NSERC). P.E.H.M. was supported by the following scholarships: Bourse de recrutement and Bourse d’exemption from the Department of Neurosciences at the Université de Montréal and Bourse d’exemption from the Quebec ministry of higher education. M.T. holds a FRQS Junior 2 salary award and funding from the Fondation Courtois.

## Supplementary information

**Supplementary figure 1 & 2 – Related to figure 3**

**Supplementary figure 3 – Related to figure 4 & 5**

**Supplementary table 1 – Detailed diet composition**

**Supplementary table 2 – Related to figure 6**

**Supplementary table 3 – Related to figure 3**

**Supplementary table 4 – Excel file - related to figure 4**

**Supplementary table 5 & 6 – Excel file - related to figure 5**

**Supplementary table 7**

**Supplementary table 8 & 9 – Related to figure 6**

---

**Supplementary figure 1:**
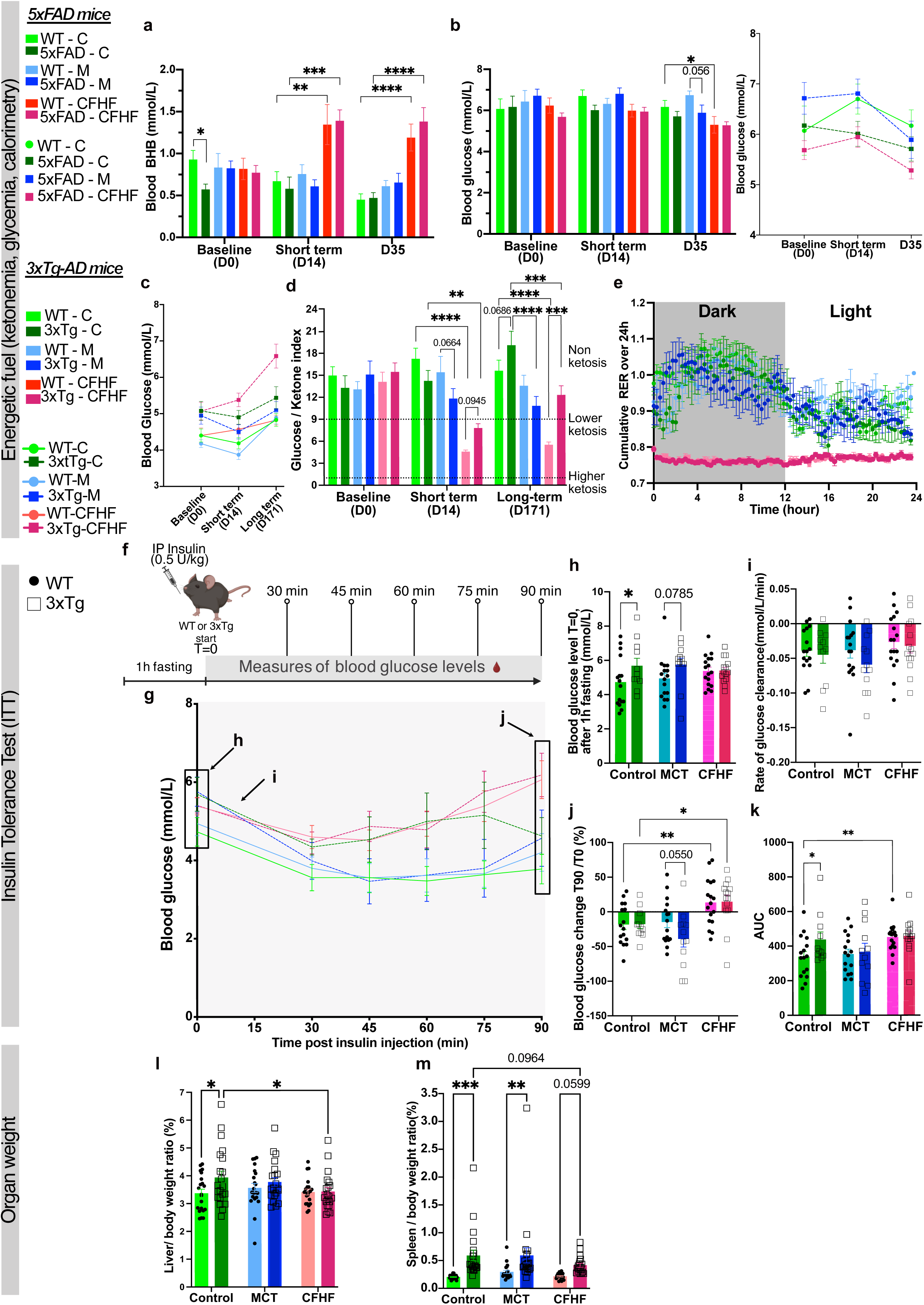
Physiological response to ketogenic interventions in 5xFAD mice and 3xTg-AD mice. a-e) Energetic fuel in : 5xFAD mice (a-b), a) Longitudinal measures of blood beta-hydroxybutyrate (BHB) (n=6-11/strain/diet, 2-way ANOVA for each timepoint (<u>Baseline</u>: not significant, <u>Short-term</u>: significant diet factor (p-value<0.0001, F(2,56)=16.15), <u>D35:</u> significant diet factor (p-value<0.0001, F(2,56)=27.38). Significant Fisher post hoc test (detailed on graph)) / b)Measures blood glucose levels (n=6-11/strain/diet, 2-way ANOVA for each timepoint (<u>Baseline and short term</u>: not significant, <u>D35:</u> significant diet factor (p-value=0.0046, F(2,56)=5.944). Significant Fisher post hoc test (detailed on graph)) and curves showing blood glucose evolution during KI in WT mice on Control diet , 5xFAD mice on Control, MCT and CFHF diets (WT-Control, 5xFAD-M, 5xFAD-CFHF) (Mixed effects model show significant time (p-value=0.0143, F(1.958, 60.69)=4.602) and experimental group factors (p-value=0.0117, F(3,38)=4.192)) in 3xTg-AD mice (c-e), c) Curves showing blood glucose evolution during KI (Mixed effects model show significant time (p-value<0.0001, F(1.988,224.7)=22.42) and experimental group factors (p-value<0.0001, F(5,114)=16.71) with interaction nearly significant (p-value=0.0513, F(10,226)=1.864)/ d)Glucose ketone index is the ratio of blood glucose levels out of blood ketones (GKI<10 indicate ketosis state) (2-way ANOVA for each timepoint (<u>Baseline</u>: not significant, <u>Short-term</u>: significant diet factor (p-value<0.0001, F(2,115)=27.42) and significant interaction between diet and strain factors (p-value=0.0235, F(2,115)=3.876), <u>Long-term</u>: significant diet factor (p-value<0.0001, F(2,112)=20.89) and strain factor (p-value=0.0212, F(1,112)=5.467) with interaction (p-value=0.0016, F(2,112)=6.856). Significant Fisher post hoc test (detailed on graph)) e) Cumulative measures of respiratory exchange ratio(RER) during 24h showing variation from the dark phase to the light phase (n=5-7/strain/diet, Mixed effects model showing significant time (p-value<0.0001, F(5.937, 198.4)=8.415) and experimental factors (p-value<0.0001, F(5,29)=18.96)). f-k) Intra peritoneal insulin tolerance test (IPITT) after 4 months on diet (n=11-17/strain/diet) showing g)Curves of blood glucose change after IP injection of insulin ITT results (Mixed effects model showing significant time (p-value<0.0001, F(3.202, 254.9)=22.36) and experimental factors (p-value=0.0119, F(5,80)=3.155) with interaction (p-value<0.0001, F(25,398)=2.571)) , h)fasting glucose levels (2-way ANOVA showing significant strain factor (p-value=0.0249, F(1,80)=5.226). Significant Fisher post hoc test (detailed on graph)); i)Rate of glucose clearance 30 min after injection (2-way ANOVA non significant); j)Rate of glucose change at T90 (2-way ANOVA showing significant diet factor (p-value<0.0001, F(2,80)=12.62). Significant Fisher post hoc test (detailed on graph)); k)Area under the curve (AUC, 2-way ANOVA showing significant diet factor (p-value=0.0114, F(2,80)=4.735). Significant Fisher post hoc test (detailed on graph)) l-m) Peripheral organ weight. / l) Liver-weight-to-body-weight ratio (n=20-21/strain/diet, 2-way ANOVA showing nearly significant strain factor (p-value=0.0492, F(1,119)=3.948). Significant Fisher post hoc test (detailed on graph)). / m) Spleen-weight-to-body-weight ratio (n=19-21/strain/diet, 2-way ANOVA showing significant strain factor (p-value<0.0001, F(1,118)=24.56). Significant Fisher post hoc test (detailed on graph)) Fisher post hoc test: *p-value<0.05; **p-value<0.01; ***p-value<0.001; ****p-value<0.0001

**Supplementary figure 2:**
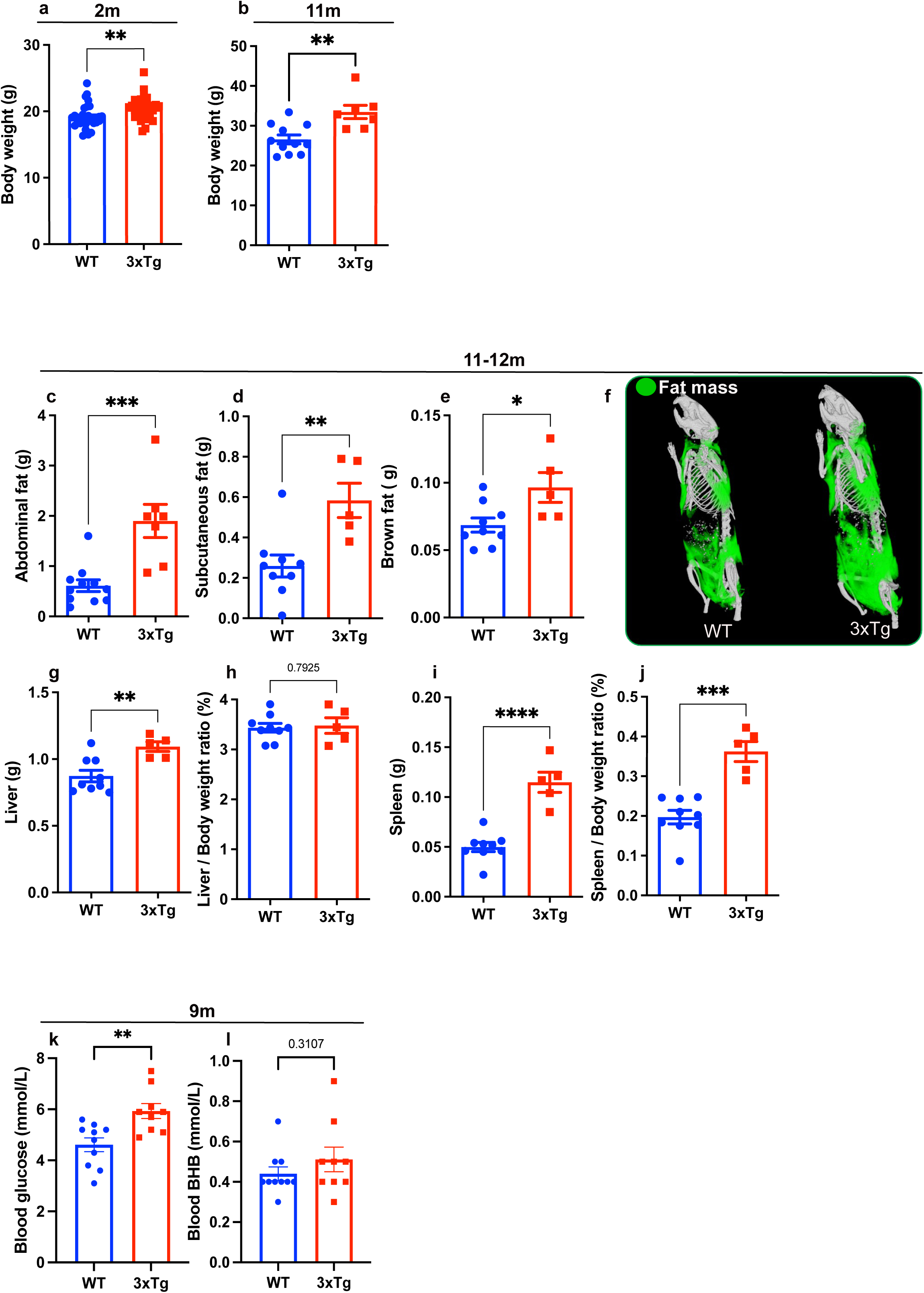
Systemic abnormalities in 3xTg-AD mice fed with regular Chow diet. a) Body weight of 2months old (2m) female mice (n=32-33/strain, Significant Unpaired two-tailed t-test: t=2.877, df=63) ; b) Body weight of 11months old (11m) female mice(n=7-11/strain, Significant Unpaired two-tailed t-test: t=3.591, df=16) ; c-e)Peripheral white and brown adipose tissue in 11m mice (n=7-11/strain): c) Abdominal fat (Significant Unpaired two-tailed t-test: t=4.317, df=16) d)Subcutaneous fat (Significant Unpaired two-tailed t-test: t=3.386, df=12), e)Brown back fat (Significant Unpaired two-tailed t-test: t=2.603, df=12) f) Representation of global body fat mass distribution in 12months old mice (12m) observed by micro-computed tomography ; g-j)Weight and weight-to-body-weight ratio of spleen and liver in 11m mice (n=5-11/strain) ; g) Liver weight (Significant Unpaired two-tailed t-test: t=3.421, df=12), i)Spleen weight (Significant Unpaired two-tailed t-test: t=6.704, df=12), j)Spleen-weight-to-body-weight ratio (Significant Unpaired two-tailed t-test: t=5.611, df=12) k-l)Circulating levels of glucose and beta-hydroxybutyrate(BHB) in 9months old mice (9m, n=9-10/strain); k) Glucose (Significant Unpaired two-tailed t-test: t=3.315, df=17) Stats: Unpaired two-tailed t-test, mean with SEM, significance: *p-value<0.05; **p-value<0.01; ***p-value<0.001; ****p-value<0.0001

**Supplementary figure 3:**
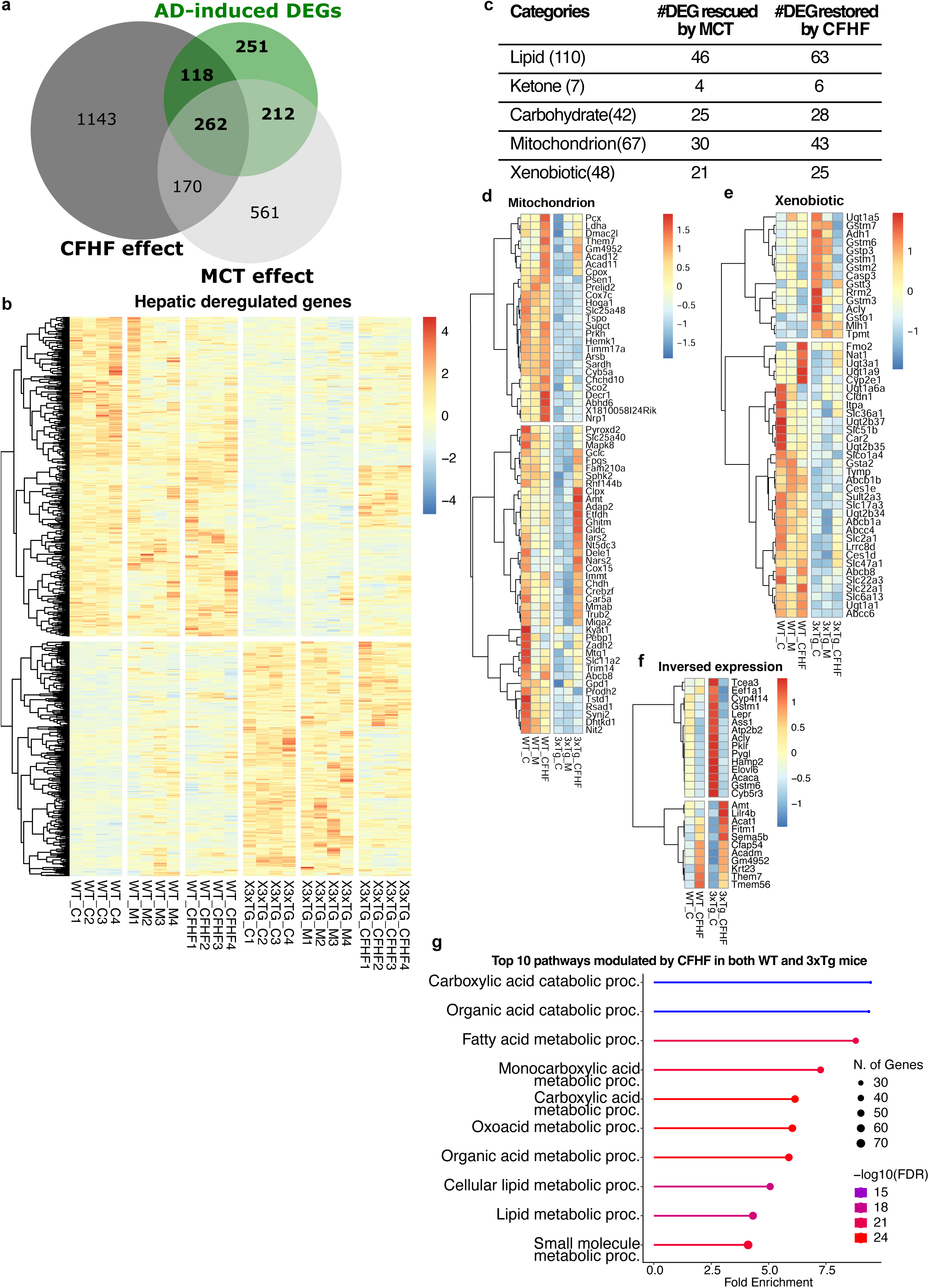
Dysregulated genes in 3xTg-AD mice liver are involved in several metabolic pathways. a) Venn diagram comparing the following contrasts : AD-induced DEGs (WT-C vs 3xTg-C), CFHF effect ( WT-C vs 3xTg-CFHF), MCT effect (WT-C vs 3xTg-M) b) Heatmap of 843 DEGs between WTC and 3xTgC (n=4/strain/diet, DESeq2, cut-off p-value<0.01) c) DEGs regrouped in genes categories and proportion rescued by MCT or CFHF d) Heatmap of Mitochondrial genes, e) Heatmap of genes involved in Xenobiotic pathways , f) Heatmap of genes inversed from Control diet to CFHF in 3xTg-AD mice g) Top 10 pathways associated with genes modulated by CFHF in WT and 3xTG

**Supplementary Table 1 :**
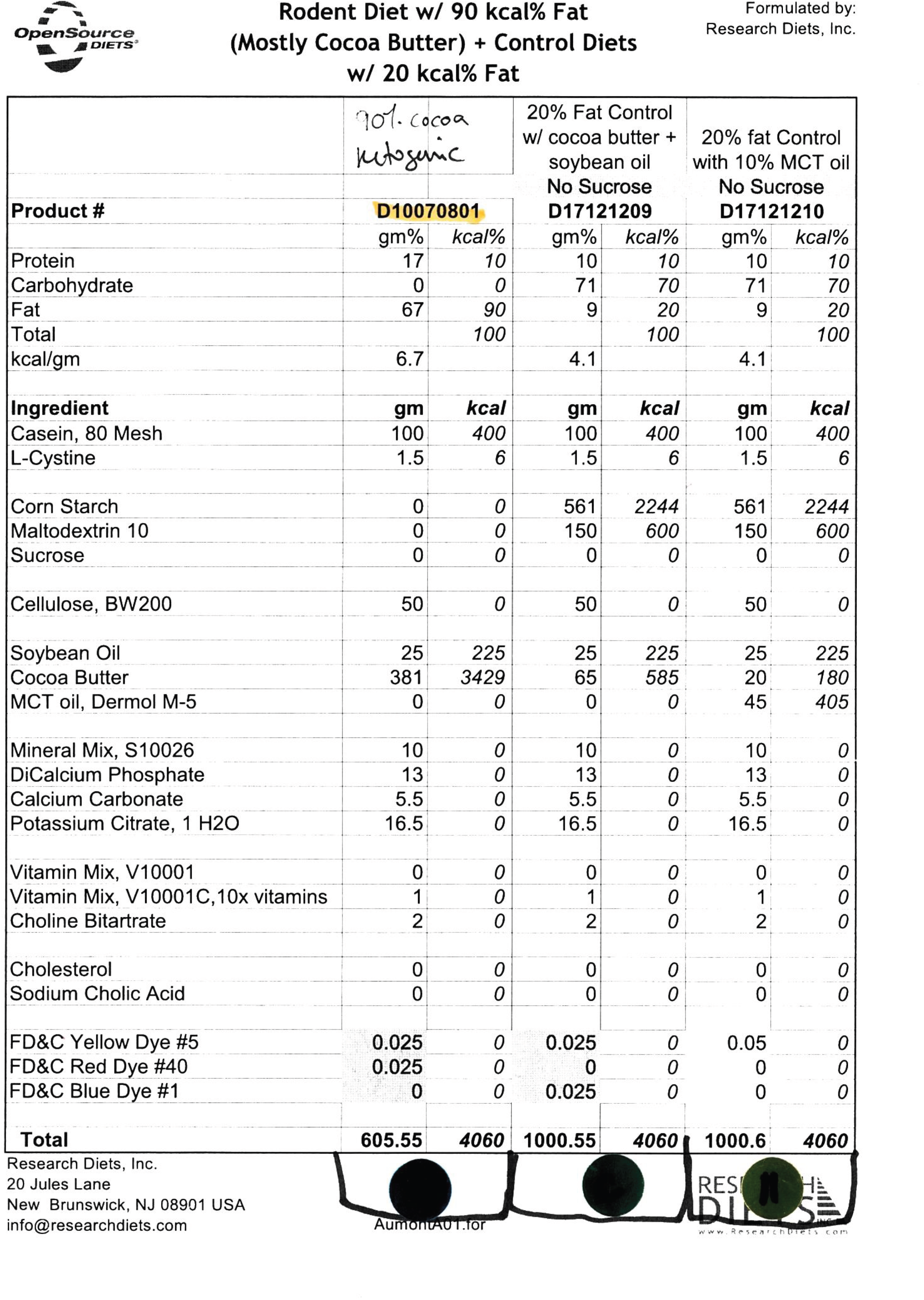
Detailed dietary composition.

|  |  |  |  |  |  |  |
| --- | --- | --- | --- | --- | --- | --- |
|  | *90% cocoa*<br>*ketogenic* |  | 20% Fat Control<br>w/ cocoa butter +<br>soybean oil<br>No Sucrose<br>**D17121209** |  | 20% fat Control<br>with 10% MCT oil<br>No Sucrose<br>**D17121210** |  |
| **Product #** | **D10070801** |  |  |  |  |  |
|  | gm% | kcal% | gm% | kcal% | gm% | kcal% |
| Protein | 17 | 10 | 10 | 10 | 10 | 10 |
| Carbohydrate | 0 | 0 | 71 | 70 | 71 | 70 |
| Fat | 67 | 90 | 9 | 20 | 9 | 20 |
| Total |  | 100 |  | 100 |  | 100 |
| kcal/gm | 6.7 |  | 4.1 |  | 4.1 |  |
| **Ingredient** | **gm** | **kcal** | **gm** | **kcal** | **gm** | **kcal** |
| Casein, 80 Mesh | 100 | 400 | 100 | 400 | 100 | 400 |
| L-Cystine | 1.5 | 6 | 1.5 | 6 | 1.5 | 6 |
| Corn Starch | 0 | 0 | 561 | 2244 | 561 | 2244 |
| Maltodextrin 10 | 0 | 0 | 150 | 600 | 150 | 600 |
| Sucrose | 0 | 0 | 0 | 0 | 0 | 0 |
| Cellulose, BW200 | 50 | 0 | 50 | 0 | 50 | 0 |
| Soybean Oil | 25 | 225 | 25 | 225 | 25 | 225 |
| Cocoa Butter | 381 | 3429 | 65 | 585 | 20 | 180 |
| MCT oil, Dermol M-5 | 0 | 0 | 0 | 0 | 45 | 405 |
| Mineral Mix, S10026 | 10 | 0 | 10 | 0 | 10 | 0 |
| DiCalcium Phosphate | 13 | 0 | 13 | 0 | 13 | 0 |
| Calcium Carbonate | 5.5 | 0 | 5.5 | 0 | 5.5 | 0 |
| Potassium Citrate, 1 H<sub>2</sub>O | 16.5 | 0 | 16.5 | 0 | 16.5 | 0 |
| Vitamin Mix, V10001 | 0 | 0 | 0 | 0 | 0 | 0 |
| Vitamin Mix, V10001C, 10x vitamins | 1 | 0 | 1 | 0 | 1 | 0 |
| Choline Bitartrate | 2 | 0 | 2 | 0 | 2 | 0 |
| Cholesterol | 0 | 0 | 0 | 0 | 0 | 0 |
| Sodium Cholic Acid | 0 | 0 | 0 | 0 | 0 | 0 |
| FD&C Yellow Dye #5 | 0.025 | 0 | 0.025 | 0 | 0.05 | 0 |
| FD&C Red Dye #40 | 0.025 | 0 | 0 | 0 | 0 | 0 |
| FD&C Blue Dye #1 | 0 | 0 | 0.025 | 0 | 0 | 0 |
| **Total** | **605.55** | **4060** | **1000.55** | **4060** | **1000.6** | **4060** |

Amount for
RESEARCH DIETS www.researchdiets.com

**Supplementary Table 2.**
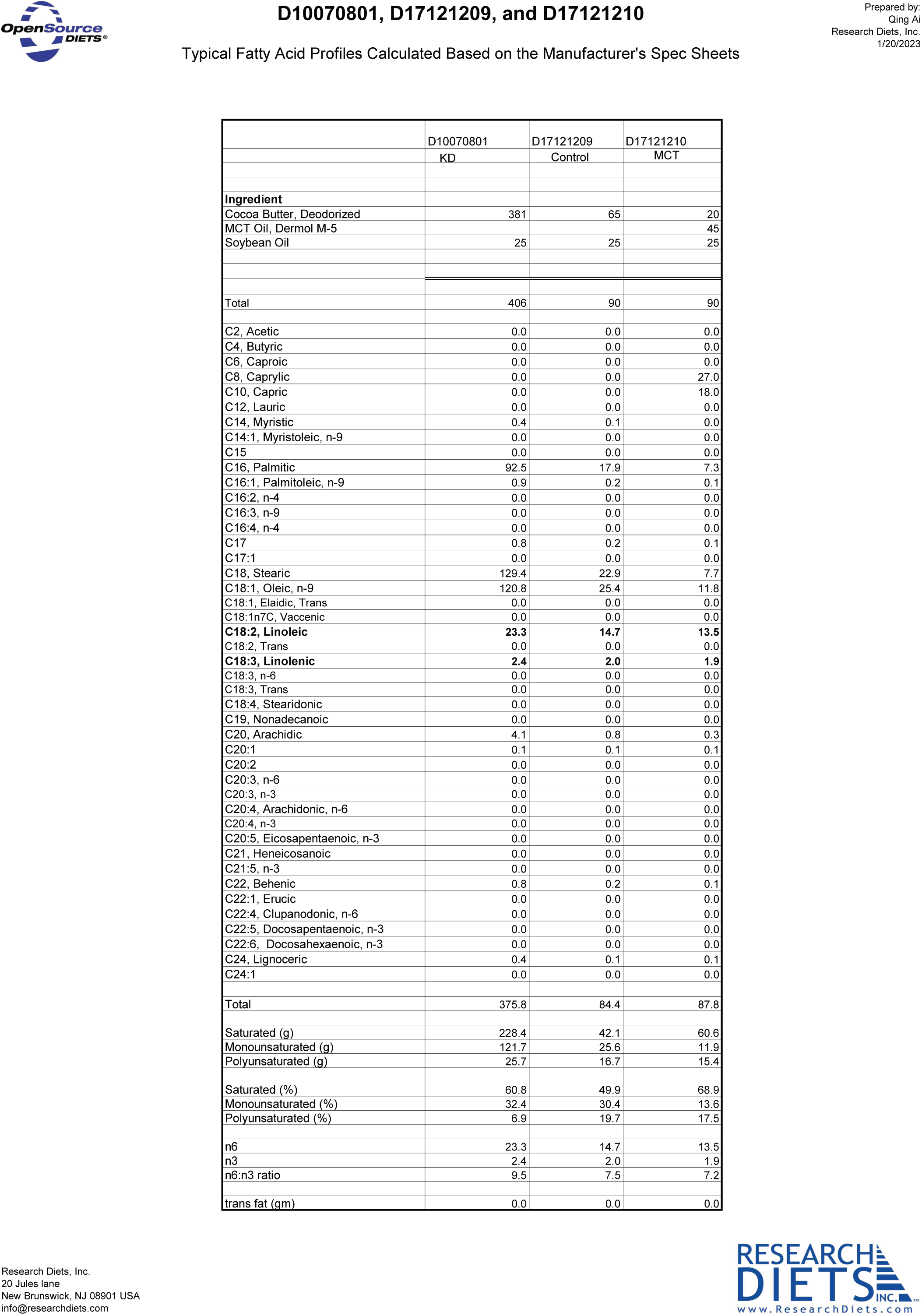

**Supplementary Table 3.**
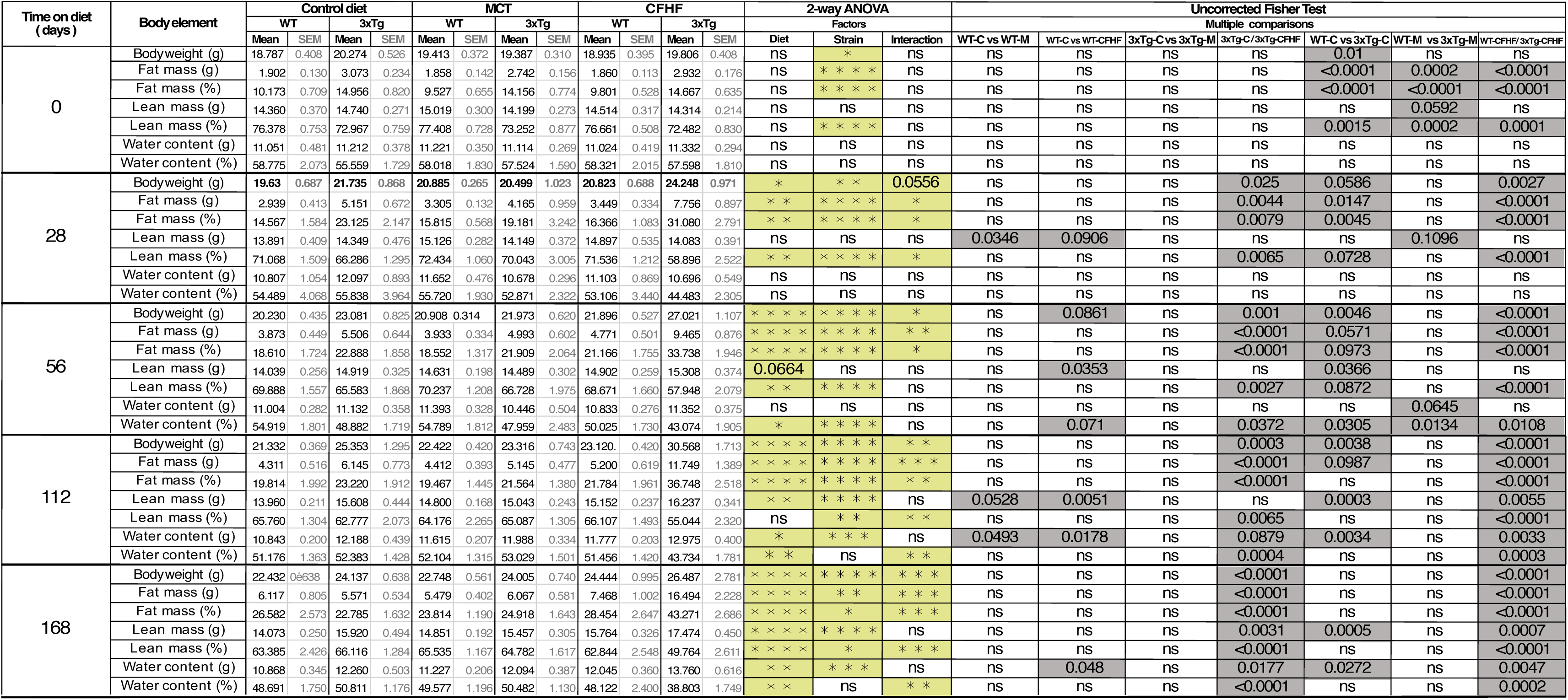
Body composition analysis giving the body weight, lean mass, fat mass and water content, expressed in grams and percentage of body weight.

**Supplementary table 7 :** Plasma levels of metabolic markers.

| Proteins | Plasmatic levels of proteins (pg/mL) |  |  |  |  |  | 2-way ANOVA |  |  | Uncorrected Fisher Test |  |  |  |  |  |  |
| --- | --- | --- | --- | --- | --- | --- | --- | --- | --- | --- | --- | --- | --- | --- | --- | --- |
|  | Control diet |  | MCT |  | CFHF |  | Factors |  |  | Multiple comparisons |  |  |  |  |  |  |
|  | WT | 3xTg | WT | 3xTg | WT | 3xTg | Diet | Strain | Interaction | WT-C vs WT-M | WT-C / WT-CFHF | 3xTg-C vs 3xTg-M | 3xTg-C / 3xTg-CFHF | WT-C vs 3xTg-C | WT-M vs 3xTg-M | WT-CFHF / 3xTg-CFHF |
| Ghrelin | 24189.824±8978.941 | 32360.830±6684.422 | 13073.385±1053.956 | 42166.410±10786.395 | 20673.399±3528.755 | 20694.082±3532.358 | ns | 0.0341 | ns | ns | ns | ns | ns | ns | * * | ns |
| Gastric inhibitory polypeptide (GIP) | 3075.305±694.640 | 3070.904±100.046 | 3179.560±324.045 | 4584.295±1163.640 | 2922.272±241.912 | 3034.760±186.438 | ns | ns | ns | ns | ns | ns | ns | ns | ns | ns |
| Glucagon-like peptide 1 (GLP-1) | 372.270±47.959 | 616.393±212.057 | 219.940 ±114.931 | 676.081 ±245.062 | 270.378±44.434 | 410.490±74.970 | ns | ns | ns | ns | ns | ns | ns | ns | ns | ns |
| Insulin | 5247.596±1258.020 | 13832.713±7742.593 | 4087.288 ±1745.046 | 13937.493±6240.077 | 2540.290 ±278.999 | 5714.642 ±1079.140 | ns | ns | ns | ns | ns | ns | ns | ns | ns | ns |
| Leptin | 4154.263±537.341 | 4619.579±575.156 | 3396.558±590.559 | 6266.293±1300.640 | 4281.879±1096.191 | 12124.620±1629.858 | 0.0011 | <0.0001 | 0.0028 | ns | ns | ns | * * * * | ns | 0.0571 | * * * * |
| Plasminogen activator inhibitor-1 (PAI-1) | 1004.327±307.319 | 2053.773±336.793 | 946.0±219.030 | 1762.344±208.10 | 1268.779±308.078 | 2361.397±187.715 | ns | 0.0001 | ns | ns | ns | ns | ns | * * | * | * * |
| Resistin | 116604.491 ±17307.271 | 141545.659±35258.750 | 138903.977±13488.080 | 150132.746±24253.772 | 182780 ±17590.293 | 114174.758±8595.101 | ns | ns | ns | ns | * | ns | ns | ns | ns | * |
| Glucagon | 3404.073±783.578 | 6277.190±2011.688 | 1540.677 ±441.962 | 6793.222±3231.706 | 1643.693±288.603 | 4462.408±780.877 | ns | 0.0124 | ns | ns | ns | ns | ns | ns | * | ns |

**Supplementary table 8 :**
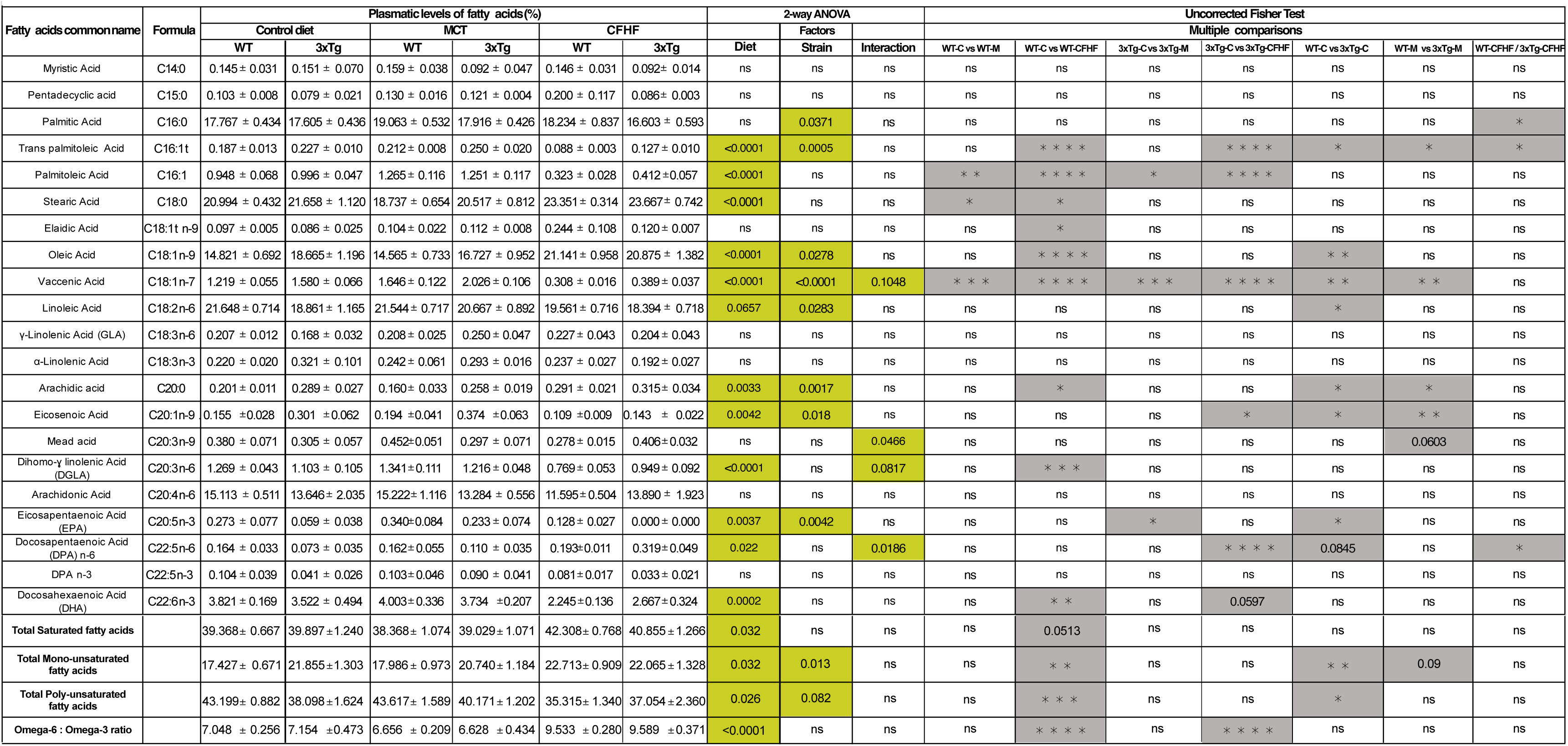
Plasma relative concentrations (%) of fatty acids identified by GC-FID.

**Supplementary table 9 :**
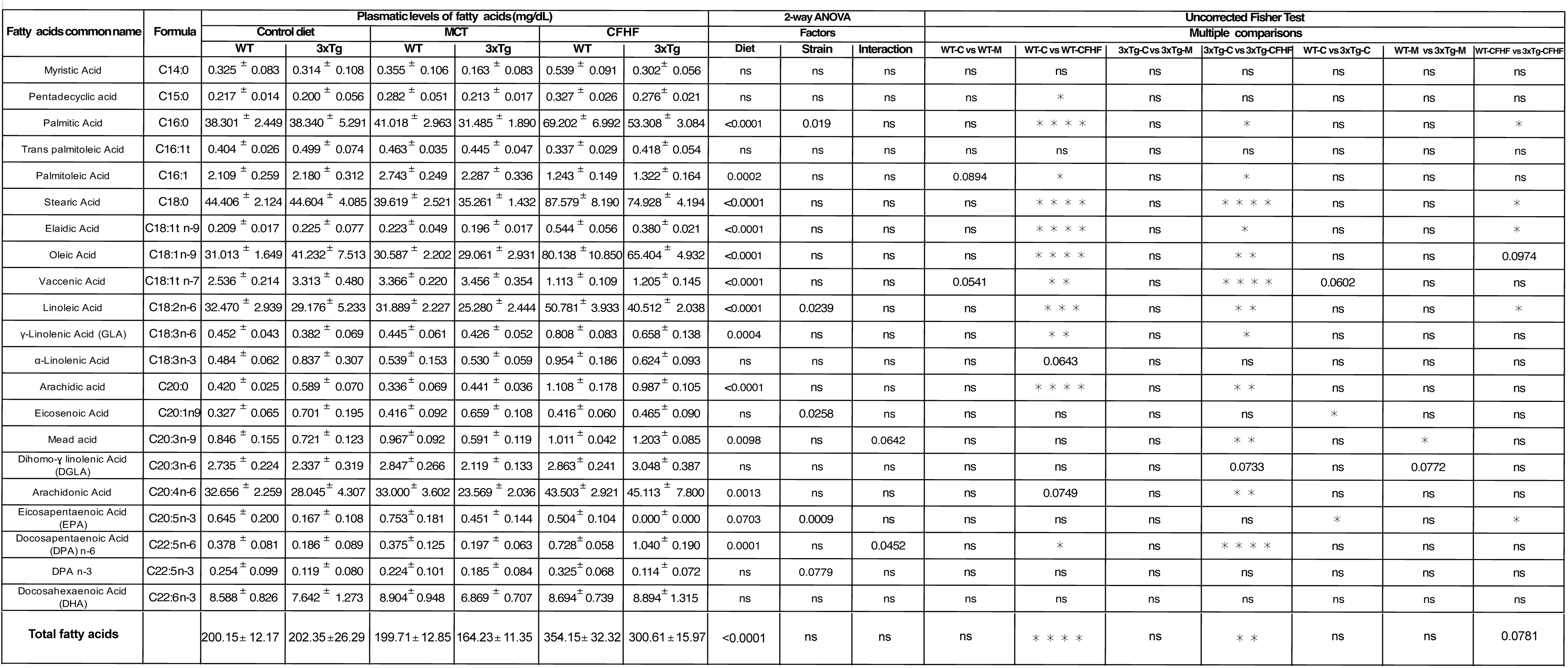
Plasma concentrations (mg§dl) of fatty acids identified by GC-FID.

